# The opposing mechanisms by which miRNAs critically contribute to differential roles of Nrf1 and Nrf2 in modulating the epithelial-mesenchymal transformation of hepatocellular carcinoma

**DOI:** 10.1101/2024.11.30.626196

**Authors:** Juan Chen, Jing Feng, Yuping Zhu, Shaofan Hu, Yiguo Zhang

**Author notes:** Contributed equally to this work (Juan Chen, Jing Feng).

## Abstract

Accumulation of various genetics and epigenetics alterations are accepted to result in the initiation and progression of hepatocellular carcinoma (HCC), and its high metastasis is viewed as a critical bottleneck leading to its treatment failure. Amongst them, the abnormal expression of several microRNAs arising from lack of antioxidant transcription factor Nrf2 (encoded by *Nfe2l2*) leads to cancer metastasis. However, much less known is about regulation of microRNAs by Nrf1 (encoded by *Nfe2l1*), even though it acts as an essential determinon of cell homeostasis by governing the transcriptional expression of those driver genes contributing to the epithelial–mesenchymal transition (EMT) involved in its metastasis. In this study, distinctive EMT phenotypes were unveiled to result from specific knockout of Nrf1 and Nrf2 in HepG2 cells, as accompanied by differential migratory and invasive capabilities. The *Nrf1α*^*−*/*−*^-leading EMT results from a significantly decrease in the epithelial *CDH1* expression, plus another increased expression of the mesenchymal *CDH2*. Such distinct phenotypes of *Nrf1α*^*−*/*−*^ from *Nrf2*^*−*/*−*^ cell lines were also attributable to differential regulation of two key microRNAs, i.e., *miR3187-3P* and *miR1247-5p*. Further experiments unraveled that Nrf1 activates the expression of *miR-3187-3p*, directly targeting for the inhibition of *SNAI1*, leading to CDH1 activation but CDH2 inhibition insomuch as to prevent the process of EMT. By contrast, Nrf2 inhibits the expression of *miR1247-5p*, relieving its inhibitory effect on MMP15 and MMP17 to promote the EMT. Collectively, these demonstrate that the EMT of liver cancer cells is likely prevented by Nrf1 *via* the miR3187-3P signaling to the SNAI1-CDH1/2 axis, but conversely promoted by Nrf2 through the miR1247-5p-MMP15/17 signaling axis.

## 1. Introduction

According to the statistic evaluation by the International Agency for Research on Cancer (IARC) of the World Health Organization (WHO), the morbidity of liver cancer is ranked world-widely in the sixth place of all new cancer cases, with its mortality rate being reached to the third[1]. Only in China, the number of newly-increased cases with liver cancer has highly reached to 410,000, at the fifth ranked in the incidence of all various types of cancer. The number of deaths from liver cancer has reached to 390,000; that is, its mortality rate is at the second highest in China. Such refractory liver cancer has heretofore been recognized as one of the main factors endangering human health [1, 2], because the detailed mechanisms underlying its malignant progression and metastasis remain elusive.

As one key issue in liver cancer treatment, the high invasiveness and metastasis of tumor cells is widely accepted to be attributable to the increasing epithelial–mesenchymal transition (EMT) in the process, albeit the EMT has been, for many years, described mainly in the embryonic development stage [3]. Specially, the occurrence and progression of liver cancer are modulated predominantly by EMT-related transcription factors, including canonical SNAI11, SNAI2, ZEB1/2, Twist, and other non-canonical such as WNT1/β-catenin, Prrx1, c-Myc and TTF1 [4-8]. Besides, the EMT process is also activated by a variety of signaling pathways such as TGF-β, MAPK, PI3K, Akt, and PTEN, controlling the expression of downstream genes responsible for tumor metastasis, but depending on the inhibitory effect of the above transcription factors in different biological backgrounds [9]. The regulatory pattern of those non-targeted transcription factors has been gradually reported by EMT markers CDH1, CDH2 and VIM, as well as other non-labeled proteins [10-12]. During EMT, inhibition of the E-cadherin (CDH1, its expression resulting in typically polygonal, cobblestone-like epithelial shapes) can enable it to acquire another spindle mesenchymal morphology, along with the expression of mesenchymal-related markers, e.g., N-cadherin (CDH2), vimentin and fibronectin [13]. In the meantime, cell junctions (tight, adhesion, gap junctions and desmosomes) were gradually disintegrated, such that cell polarity was lost, actin expression was increased, pseudopodia were formed, cell adhesion was promoted, and the expression of N-cadherin and integrin was enhanced, but the epithelial cell adhesion was reduced. This is accompanied by differential expression of matrix metallopeptideases (MMPs, which is a family of zinc- and calcium-dependent proteolytic enzymes to degrade almost all components of the extracellular matrix (ECM)) [14].

Intriguingly, the abnormal expression of microRNAs (miRNAs) in tumor cells has clearly been shown to promote the occurrence, development, migration and invasion of cancer cells in many ways compared with normal tissues [4, 10, 15, 16]. That is, the abnormal miRNA expression is accompanied by the development of carcinogenic transformation, i.e., EMT, with the high metastasis and invasion ability [17]. Of note, the EMT of tumor cells were reportedly regulated by a class of miRNAs, including miR29b, miR30a, miR30c, miR137, miR34a, and miR15, through transcription factor SNAI1 [18-23], exerting distinct biological effects of tumor suppressors or promoters in different types of cancers [24]. Besides, the EMT of tumor cells is also regulated by nuclear factor erythroid 2-related factor 2 (Nrf2, encoded by *Nfe2l2*) through distinct miRNA pathways. However, less known is about how Nrf2, together with nuclear factor erythroid 2-related factor 1 (Nrf1, encoded by *Nfe2l1*), regulates those critical miRNAs for differentially controlling the development of EMT in liver cancer cells.

In mammalians, Nrf1 and Nrf2 are two principal members of the cap’n’collar basic region-leucine zipper (CNC-bZIP) transcription factor family that governs cellular (redox, energy, metabolism) homeostasis and organ integrity [25]. Indeed, prior studies have clearly shown that Nrf1 and Nrf2 interact with each other in the regulation of redox, glucose, and lipid metabolisms and malignant proliferation of hepatocellular carcinoma [26]. However, it has been only reported that Nrf2 activation can promote the occurrence and development of EMT, and the persistent activation of Nrf2-mediated miRNA regulation of EMT in cancer cells has been also widely reported [27]. In this study, we discovered that specific knockout of Nrf1 and Nrf2 in HepG2 cells results in distinct EMT phenotypes, as accompanied by differential migratory and invasive capabilities. Such distinct phenotypes of *Nrf1α*^*−*/*−*^ from *Nrf2*^*−*/*−*^ cell lines were also attributable to differential regulation of two key miRNAs, i.e., *miR3187-3P* and *miR1247-5p*. Further evidence has been provided revealing that Nrf1*α* activates the expression of *miR-3187-3p*, directly targeting for inhibition of *SNAI1*, thereby leading to CDH1 activation, but CDH2 inhibition, insomuch as to prevent the EMT process. By contrast, Nrf2 inhibits *miR1247-5p*, relieving its inhibitory effects on MMP15 and MMP17 to promote EMT. Collectively, these demonstrate that the EMT of liver cancer cells is much likely prevented by Nrf1*α via* miR3187-3P signaling to the SNAI1-CDH1/2 axis, but rather promoted by Nrf2 *via* the miR1247-5p-MMP15/17 signaling axis.

## 2. Materials and methods

### 2.1. Cell lines, culture, and transfection

The *Nrf1 α*^*−/−*^ cells used in this study were constructed by TALENs-mediated genome editing of HepG2 cells, while *Nrf2*^*−*/*−*^ cells were constructed by CRISPR/Cas9-editing system through HepG2 cells [28]. These cell lines were cultured in DMEM media containing 5 mM glutamine, 10 % (v/v) of fetal bovine serum (FBS) and 100 units/mL of penicillin and streptomycin at 37 °C in a 5 % CO_2_ incubator. The experimental cells were transfected with indicated plasmids-contained Lipofectamine® 3000 reagent and then cultured for 8 h in Opti-MEM (Gibco, Walsam, MA, USA). These cells were allowed for 24 h recovery from transfection in a fresh complete medium before the following experiments were performed.

### 2.2. Expression constructs for Nrf1, Nrf2 and indicated reporters

Two expression constructs for human Nrf1 and Nrf2 were made by inserting their full-length cDNA sequences, respectively, into the KpnI/XbaI site of pcDNA3.1/V5His B. The seed sequences of *miR3187-3p* and *miR1247-5p* were respectively constructed into the AgeI/EcoRI site of PLKO.1-TRC cloning vector. Those indicated *3×ARE s*ites of *miR3187-3p* (i.e., 1# to 5#) and *miR1247-5p* (i.e., 1# to7#) were synthesized and ligated into the KnpI/XhoI site of PGL3-Promoter vector. The resulting *ARE-Luc* reporters were hence created by inserting the consensus *ARE*-adjoining sequences from indicated gene promoters. Lastly, the 3’-UTR sequences of *SNAI1, MMP15, MMP17*, and *MMP25* were also subcloned into the XhoI/NotI site of psiCHECK-2 plasmid, respectively.

### 2.3. Real-time qPCR analysis of mRNA expression levels

Total RNAs were extracted from experimental cells by using an RNA extraction kit (TIANGEN, Beijing, China), and then approximately 2.0∼2.5 μg of total RNAs were added in a reverse-transcriptase reaction to generate the first strand of cDNA (by using the Revert Aid First Strand Synthesis Kit, Thermo, Waltham, MA, USA). The synthesized cDNA served as the template for quantitative PCR (qPCR) in the GoTaq®qPCR Master Mix (Promega, Madison, WI, USA). Subsequently, the mRNA expression levels were measured by RT-qPCR with indicated pairs of primers (as listed in Table S1). The mRNA expression level of β-actin served as an optimal internal standard control and all relative mRNA expression abundances of other genes were hence presented as fold changes. Of note, the expression levels of miRNA need to involve specific reverse transcription primers to ensure its specificity (as shown in Table S2).

### 2.4. Western blotting analysis of protein expression abundances

Experimental cells were harvested in a denatured lysis buffer (0.5% SDS, 0.04 mol/L DTT, pH 7.5, containing 1 tablet of complete protease inhibitor EASYpacks in 10 ml of this buffer). The total lysates were further denatured by boiling at 100 °C for 10∼15 min, sonicated sufficiently, and diluted with 3 × loading buffer (187.5 mmol/L Tris-HCl, pH 6.8, 6% SDS, 30% Glycerol, 150 mmol/L DTT, 0.3% Bromphenol Blue), before being re-boiled at 100°C for 5 min. Thereafter, equal amounts of protein extracts were subjected to separation by SDS-PAGE, and then transferred to polyvinylidene fluoride (PVDF) membranes (Millipore, Billerica, MA, USA), before being visualized by Western blotting with distinct antibodies (Table S3). β-actin served as an internal control to verify equal amounts of proteins loaded in each of the electrophoretic wells.

### 2.5. Luciferase reporter assay

Equal numbers (1.0 × 10^5^) of HepG2 cells were allowed for growth in each well of 12-well plates. After reaching 75% ∼85% confluence, the cells were co-transfected for 8 h with an indicated luciferase plasmid alone or together with one of indicated expression constructs mixed with the Lipofectamine®3000 agent in Opti-MEM (Gibco, Waltham, MA, USA), in which the *pRL-TK* reporter served as an internal control for transfection efficiency. After being recovered for 24 h in a fresh complete medium, the cells were lysed and then subjected to the dual-reporter assay (Promega, Madison, WI, USA). The ARE-driven luciferase reporter activity was calculated by normalization to the internal Renilla activity, whilst specific target gene-mediated luciferase reporter activity was evaluated by further normalization to background activity measured from an empty expression vector being transfected with ARE-driven luciferase plus *pRL-TK* reporters. The resulting data are graphically shown as Mean± S.D. of at least three independent experiments performed in triplicates.

### 2.6 The transwell-based migration and invasion assays

The transwell-based cell migration and invasion were assayed in the modified Boyden chambers (Transwell, Corning Inc. Lowell, MA, USA). Equal numbers of cells were allowed for growth in each well of 12-well plates. After reaching 70–80% confluence, they were starved for 12h in a serum-free medium. The experimental cells (1× 10^5^) were suspended in a 0.2-ml medium containing free FBS and seeded in the upper chamber of each transwell. The cell-seeded transwells were placed in each well of 24-well plates containing 1 ml of complete medium (i.e., the lower chamber), and cultured for 24 h in the incubator at 37°C with 5% CO_2_. The remaining cells in the upper chamber were removed, whilst the cells attached to the lower surface of the transwell membranes were fixed with 4% paraformaldehyde (AR10669, BOSTER) and stained with 1% crystal violet reagent (Sigma) before being counted.

### 2.7. Cell scratch assay

The cells were digested with trypsin and counted after suspension, so that 3×10^5^ cells were inoculated in per well of 6-well plate (at the bottom of which 4-6 horizontally-dividing lines per well were already sculptured with a maker pen). When the cells reached more than 95 % confluence, additional 3-5 lines were drawn along the vertical direction of the pre-maker line in the hole. The cells were washed with PBS and replaced with 2 mL serum-free DMEM. The initial images of the scratch were acquired under an inverted microscope, and the cells continued to be cultured for indicated lengths of time at 5 % CO_2_ and 37 °C. Thereafter, the scratched images were recovered to distinct extents, all of which were, in the meantime, collected by microscopy at 24, 48, 72, and 96 h in serum-free DMEM media.

### 2.8. Lentiviral packaging for overexpression of indicated miRNAs

The indicated miRNA genes were respectively subcloned into the Plko.1 TRC vector and subjected to sequencing verification. The miRNA-expressing Plko.1 TRC, psPAX2 and pMD2G plasmids were combined in a 4:3:2 ratio (total mass of 24 μg DNA) and introduced into 293T cells *via* transfection. Following the digestion of such HEK 293T cells, 7 × 10^5^ cells were inoculated into a 10-cm Petri dish, subsequently with 8 mL of a high glucose medium added. The cells were cultured overnight until the bottom of the dish was covered by 70% of the cell confluence. At this point, the medium was replaced with 4 mL of another serum-reduced medium (Opti-MEM). Following a 30-minute incubation period with the transfection reagent, the plasmid transfection was performed. Following a transfection period of 8 h, the transfection medium was replaced with another standard growth medium to allow cells for 48 h recovery, and thereafter, the medium was supplemented in accordance with the rate of cell growth. The supernatants were collected at 48, 72 and 96 h post-transfection and stored at 4 °C. The concentrated lentivirus was diluted appropriately and 50 μL of the virus suspension added to each well of the experimental cells. Following a 24-h infection period, the lentiviral resistant cells were selected with 25 mg/ml of puromycin. Subsequently, the monoclonal miRNA-expressing cells were further selected, and analyzed by Western blotting and RT-qPCR. The experimental cell lines utilized in this study are listed in Table S4.

### 2.9. Clone formation assay

About 500, 1000 or 1500 experimental cells were seeded in each well of six-well plates and then cultured in 5 % CO_2_ at 37 °C for two weeks. During this period, 1 mL of fresh complete medium containing 10 % serum was added every 3 days. Finally, the cells were fixed, stained, and photographed under an inverted microscope, before being counted the number of cell clones.

### 2.10. Flow cytometry analysis of cell cycle and apoptosis

After experimental cells (3 × 10^5^) were allowed for growth in each well of 6-well plates, they were then pelleted by centrifuging at 1000×g for 5 min and washed with PBS for three times, before being incubated for 15 min with 5 μL of Annexin V-FITC and 10 μL of propidium iodide (PI) in 195 μL of the binding buffer prior to flow cytometry analysis. The resulting data were analyzed by the FlowJo 7.6.1 software (FlowJo, Ashland,OR, USA) and shown graphically. For cell cycle analysis by flow cytometry, the cells growing in the logarithmic phase were digested, centrifuged at 1000 rpm for 5 min, washed 2-3 times with PBS after the supernatants were removed. The cell pellets were resuspended in 300 uL pre-cooled PBS, followed by slow addition of 700 uL anhydrous ethanol while slightly oscillating and mixing, before being stored at 4 °C overnight. On the next day, the cells were centrifuged at 1000 rpm for 5 min at 4 °C. The cell pellets were re-suspended in 100 uL of binding Buffer and incubated for 15 min with 5μl of PI-Staining Solution and 5μL Annexin V-FITC (in the dark) at room temperature, before being subjected to flow cytometry analysis of the amounts of DNA for different periods of experimental cells.

### 2.11 Statistical analysis

Statistical significance of changes in the reporter gene activity or other gene expression was determined using either the Student’s *t-*test or Multiple Analysis of Variations (MANOVA). The resulting data are shown as a fold change (Mean ± S.D) relative to control values, which represents at least three independent experiments that were each performed in triplicates.

## 3. Results

### 3.1. Distinctive effects of Nrf1- and Nrf2-deficiencies dictating their cellular phenotypes related to EMT

To elucidate distinct impacts of Nrf1 and Nrf2 on the EMT phenotype, here we observed that cellular morphological plasticity exhibited characteristics of EMT, especially upon knockout of *Nrf1α* from HepG2 cells, as evidenced by altered morphogenesis from oval cell contours to spindle-shaped fusiform (Figure 1A, *a1, middle*). This phenomenon closely resembled that yielded by the loss of cell-cell adhesion during the EMT transformation, leading to detachment from the basement membrane and subsequent long-distance metastasis and invasion [2, 29]. By sharp contrast, no significant differences in the cell morphology were observed following knockout of *Nrf2*, when compared to that of wild-type cells (Figure 1A, *a1, right vs left panels*).

**Figure 1.**
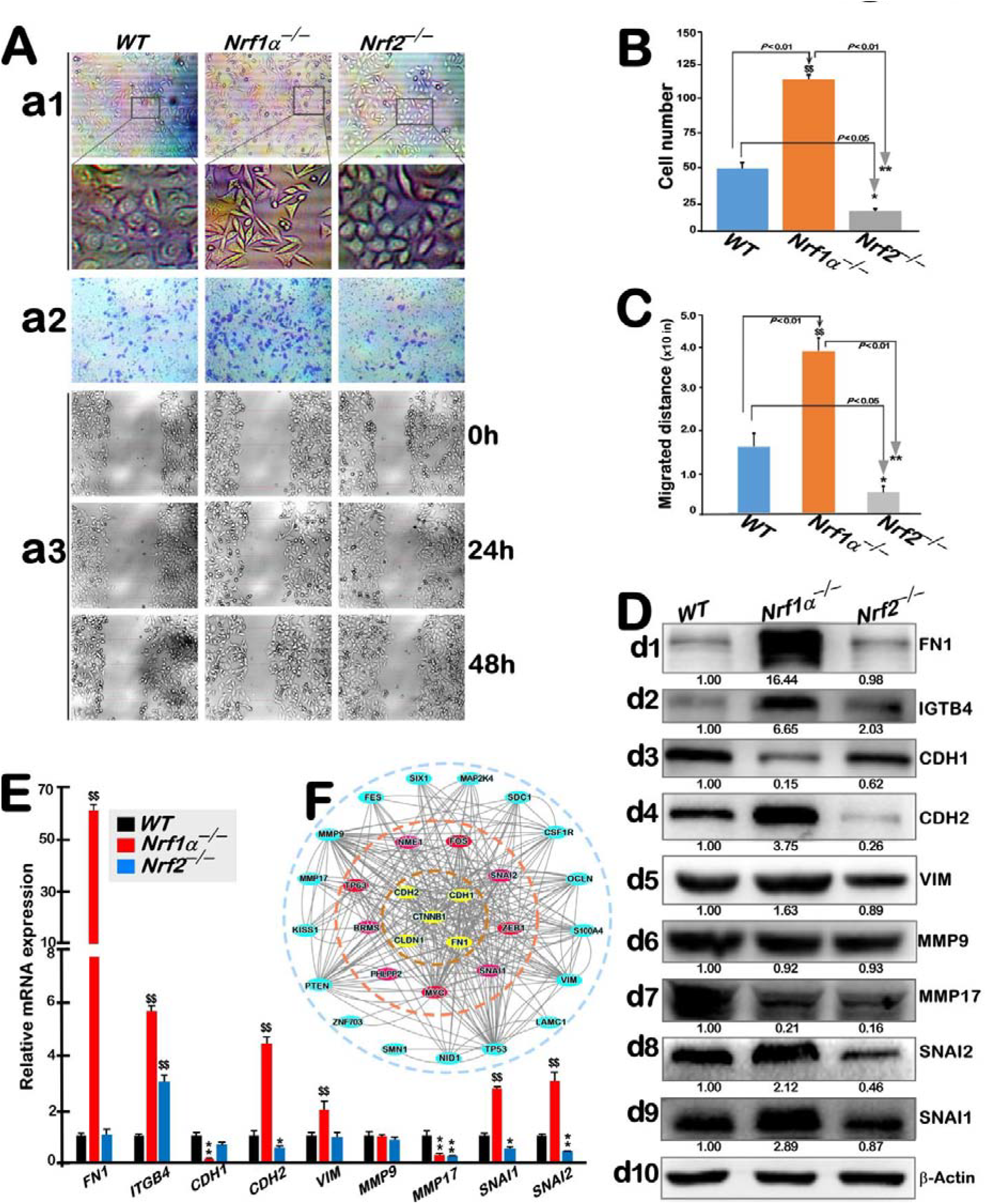
Distinct changes in the cellular morphology and relevant gene expression caused by loss of Nrf1α^*−/−*^ or Nrf2^*−/−*^. A. Distinctions in the EMT-relevant morphological plasticity of between *WT, Nrf1α*^*−/−*^, and *Nrf2*^*−/−*^ cell lines. These were evaluated by their distinctive representative images obtained from routine microscopy (*a1*), the transwell assays for their basemembrane permeability (*a2*), and another scratch assay for their migratory capacity (*a3*), respectively. B. The permeability of indicated cells throughout transwell basemembranes was subject to quantitative analysis. The results are shown graphically as Mean ± S.D. with significant increases ($$, *p* < 0.01) and significant decreases (*, *p* < 0.05; **, *p* < 0.01). All these data were determined from at least three independent experiments each performed in triplicates. C. The migratory capacity evaluated by indicated cell scratch assay was subjected to quantitative analysis. The resulting data are shown graphically as Mean ± S.D. with significant increases ($$, *p* < 0.01) and significant decreases (*, *p* < 0.05; **, *p* < 0.01). All these were determined from at least three independent experiments each performed in triplicates. D. Western blotting analysis of indicated protein expression abundances in *WT, Nrf1α*^*−/−*^, and *Nrf2*^*−/−*^ cell lines with distinct antibodies. The intensity of all immunoblots was quantified and normalized by that of β-actin, before being shown as fold changes (*on the bottom of indicated blots*). E. Real-time qPCR analysis of indicated gene expression at their mRNA levels in *WT, Nrf1α*^*−/−*^, and *Nrf2*^*−/−*^ cell lines. The resulting data are shown as fold changes (Mean ± SD), with significant increases ($$, p < 0.01), that were statistically determined from at least three independent experiments performed each in triplicates (n = 3 × 3). F. A schematic representation of the hierarchical interaction networks between the EMT-related transcription factors and target genes, which were drawn by using the Cytoscape software.

Next, the membrane permeability of examined cells was investigated by using transwell experiments. The resulting observations revealed that knockout of *Nrf1*α led to a notable increase in the cell membrane permeability (Figure 1A, *a2, middle*, and Figure 1B), whereas no discernible differences were observed by comparison of *Nrf2*^*−/−*^ cells with wild-type (Figure 1A, *a2, right vs left panels*). Subsequently, the migration capacity of examined cell types was also evaluated by a cell scratch assay. The experimental observations demonstrated that the migration distance of *Nrf1α*^*−/−*^ cells was significantly greater than that of *Nrf2*^*−/−*^ cells and even much greater than that of the control cells, remarkably occurring at 48 h (Figure 1A, *a3*). Of note, the migration of *Nrf1α*^*−/−*^ cell was incrementing as the time increases, and hence *Nrf1α*^*−/−*^ cells exhibited a greater capacity than those of both *WT* and *Nrf2*^*−/−*^ cell lines migrated at varying distances (Figure 1C).

Subsequently, both protein and mRNA expression levels of those EMT-related markers were examined by Western blotting and RT-qPCR, respectively. The results unraveled that a significant reduction of CDH1, a classic marker associated with the epithelial characteristics, was determined in *Nrf1α*^*−/−*^ cells (Figure 1,D and F). Conversely, varying extents of increases in all other examined protein and mRNA expression of FN1, ITGB4, CDH2, VIM, SNAI1, and SNAI2 (all of which are linked to mesenchymal genes) were exhibited in *Nrf1α*^*−/−*^ cells (Figure 1,D and E). Notably, the average expression abundance of FN1 at protein and mRNA levels reached to certain levels of nearly 60 times higher than that observed in *WT* cells. By contrast, knockout of *Nrf2*^*−/−*^ resulted in only marginal or even no changes in the above-examined proteins and their mRNA expression levels (Figure 1, D& E). However, the expression of MMP9 was almost unaffected by specific loss of *Nrf1α*^*−/−*^ or *Nrf2*^*−/−*^, albeit down-regulated expression of MMP17 by each loss of *Nrf1α*^*−/−*^ or *Nrf2*^*−/−*^ (Figure 1, D & E).

Collectively, these results demonstrate distinct effects of *Nrf1α*^*−/−*^ from *Nrf2*^*−/−*^ on its deficient cell morphogenesis that are contributable to the EMT-relevant phenotypes. Moreover, according to the regulatory relationship of the above-examined EMT molecules, including EMT markers and relevant transcription factors, along with their effects on targeting gene expression profiling, particularly arising by the loss of *Nrf1α*^*−/−*^ or *Nrf2*^*−/−*^, such an interaction diagram between the EMT-related transcription factors and target genes was presented herein (Figure 1F).

### 3.2. Specific loss of Nrf1α^−/−^ or Nrf2^−/−^ leads to differential miRNA expression profiling critically required for EMT

To determine whether such distinct phenotypes of between *Nrf1α*^*−/−*^ and *Nrf2*^*−/−*^ cell lines are dictated or modulated by differential expression profiling of those putative miRNAs, we here performed small RNA sequencing of indicated cell lines, each with two replicates. As clearly shown in Figure S3A, those samples were controlled with a rather high quality, aside from almost negligible small differences. The resulting data were subject to bioinformatics prediction by miRanda and TargetScan, revealing the target genes of 2154 miRNAs regulated in *Nrf1α*^*−/−*^ and *Nrf2*^*−/−*^ cell lines, as a result with a high reliability of such target prediction being determined by the Venn statistics (Figure S3B). In addition, some target gene intersections between miRanda and TargetScan exist (as illustrated in Figure S3C).

To decipher the correlation of miRNAs with significant differences between distinct examined groups, we performed miRNA association analysis of their significant differences. The results revealed that only 16 of significantly differentially expressed 317 miRNAs were overlapped in the three groups (Figure S3D). Further bioinformatics analysis of overlapped genes unraveled those putative Nrf1/2-regulated ARE sites existing within the promoter regions of aforementioned 16 miRNAs (as enlisted in Figure S9). In order to gain insights into the functional distribution characteristics of differentially expressed miRNAs, we conducted a GO enrichment analysis of differential miRNAs-targeting genes in *Nrf1α*^*−/−*^ or *Nrf2*^*−/−*^ cell lines compared with *WT* cells. The results showed that differential miRNAs exhibited significant gene differences in their biological processes, cellular components and molecular functions, such as biological adhesion and cell colonization (Figure S1). Concurrently, another pathway enrichment analysis of those miRNA-target genes was also conducted using the KEGG database. Such KEGG enrichment analysis of *Nrf1α*^*−/−*^ vs *WT* cell lines revealed significant differences in target genes involved in cell growth and death, transcription and translation, signal transduction, and tumour metabolism (Figure S2A). Similarly, another pathway enrichment of *Nrf1α*^*−/−*^ vs *WT* cell lines indicated significant discrepancies in signaling pathways including intercellular receptors, liver cancer, and endoplasmic reticulum (ER) protein processing (Figure S2B) between the two cell lines. These indicators aforementioned are evidently associated with the EMT process of liver cancer cells. Further comparison of the KEGG analysis of *Nrf2*^*−/−*^ vs *WT* cell lines revealed significant discrepancies in amino acid metabolism, lipid synthesis and signal transduction (Figure S2C), while pathway enrichment analysis also unveiled that their primary discrepancies of target genes were observed in those pathways associated with breast cancer, human cytomegalovirus, and bile secretion (Figure S2D).

Upon further comparison between *Nrf1α*^*−/−*^ and *Nrf2*^*−/−*^ cell lines, 232 of significant differentially expressed miRNAs were determined, of which 188 were upregulated whilst additional 44 were downregulated (Figure 2A, *left two columns*). However, the majority of miRNAs exhibited evident downregulation by the loss of *Nrf1α*^*−/−*^ versus *WT* controls (Figure 2A, *middle two columns*), but the loss of *Nrf2*^*−/−*^ caused significant upregulation of those major miRNAs (Figure 2A, *right two columns*). Such marked differences in distinct miRNA expression were also further presented by two volcano plots created from *Nrf1α*^*−/−*^ cells (Figure 2B) and *Nrf2*^*−/−*^ cells (Figure 2C), when compared with *WT* cells.

**Figure 2.**
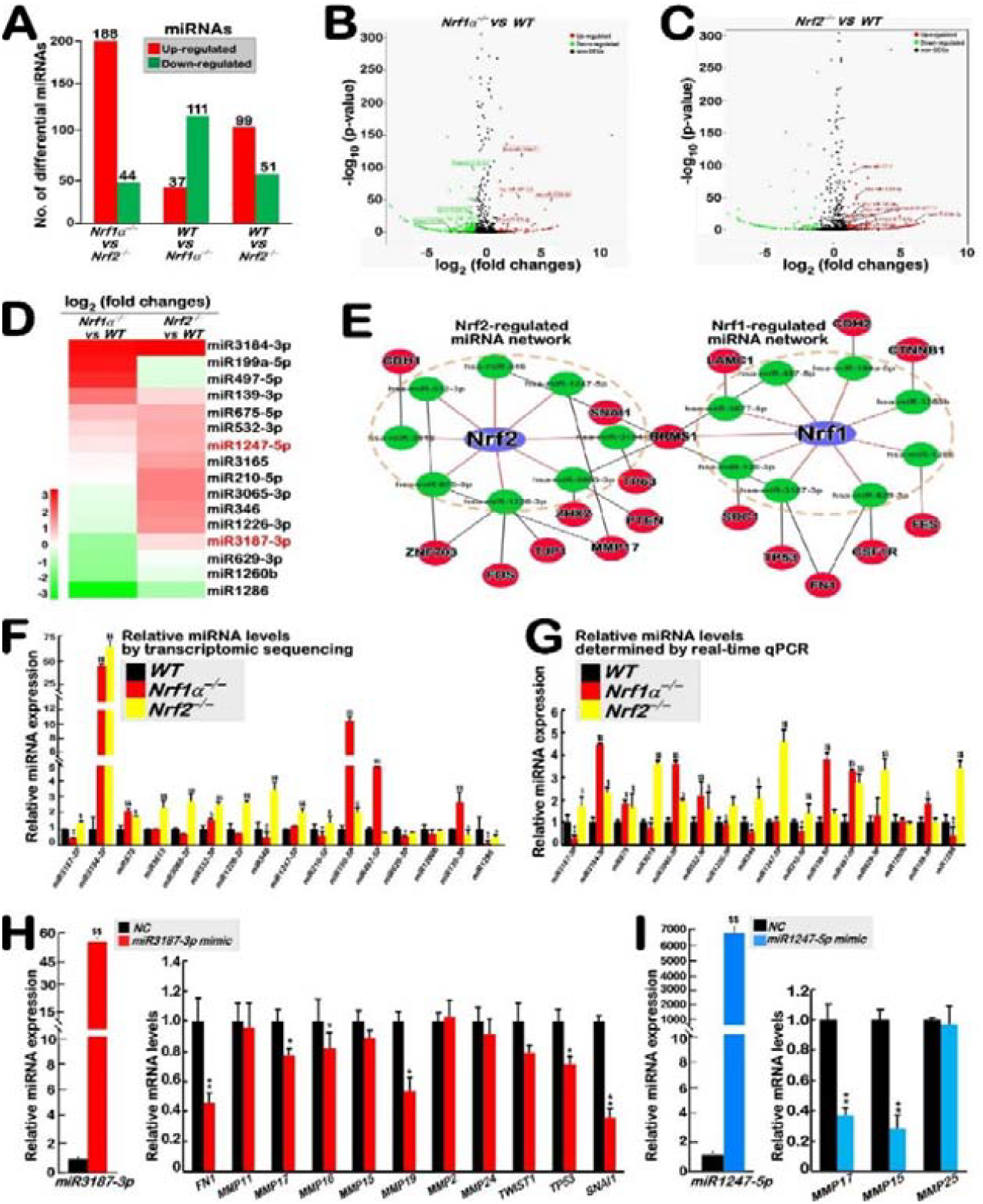
Selection of key miRNAs essentially responsible for the EMT. A. Differentially up- or down-expressed miRNAs were selected by small RNA sequencing of three examined cell lines. B,C. Two volcano plots of significant differential miRNA expression genes were created by comparison of *Nrf1α*^*−/−*^ cells (B) or *Nrf2*^*−/−*^ cells (C) with *WT* cells. Some miRNAs were marked out. D. A heatmap was made up of 16 candidate miRNAs selected from *Nrf1α*^*−/−*^ and *Nrf2*^*−/−*^ cell lines. E. Two interactive miRNA networks regulated by Nrf1 or Nrf2, along with their cognate targets, were drawn by Cytoscape. F. Relative expression levels of 16 candidate miRNAs were determined by their transcriptome sequencing. The data are shown graphically as fold changes (Mean ± S.D.) with significant increases ($, *p* < 0.05; $$, *p* < 0.01) and significant decreases (*, *p* < 0.05; **, *p* < 0.01), all of which were determined from two independent experiments each performed in triplicates. G. Relative expression abundances of the above-mentioned 16 candidate miRNAs were validated by real-time qPCR. The resulting data are shown graphically as fold changes (Mean ± S.D.) with significant increases ($, *p* < 0.05; $$, *p* < 0.01) and significant decreases (*, *p* < 0.05; **, *p* < 0.01), all of which were determined from three independent experiments each performed in triplicates. H. *Left panel* shows real-time qPCR analysis of relative miRNA expression levels of experimental *WT* cells that had been transfected with a mimic miRNA3187-3p (i.e., miR3187-3p) or another negative control (NC). *Right panel* shows relative expression levels of putative miR3187-3p-targeting downstream genes involved in EMT. The resulting data are shown graphically as fold changes (Mean ± S.D.) with significant increases ($$, *p* < 0.01) and significant decreases (*, *p* < 0.05; **, *p* < 0.01), which were determined from three independent experiments each performed in triplicates. I. Similarly real-time qPCR analysis of miR1247-5p mimic (in parallel with NC) and its downstream target gene expression was also carried out as described above.

By predicting target genes based on EMT-related markers, 8 candidate miRNAs screened from *Nrf1α*^*−/−*^ cells were miR3184-3p, miR199a-5p, miR3187-3p, miR497-5p, miR629-3p, miR1260b, miR139-3p, miR1286. Besides miR3184-3p, additional 8 miRNAs selected from *Nrf2*^*−/−*^ cells were miR675-5p, miR3165, miR3065-3p, miR532-3p, miR346, miR1226-3p, miR1247-5p, and miR210-5p. Abundances of such selected miRNA-expressing genes in *Nrf1α*^*−/−*^ or *Nrf2*^*−/−*^ versus *WT* cell lines were shown in a heatmap (Figure 2D). Two interaction diagrams of their regulatory relationships and targeting effects were inferable to be mediated by Nrf1 and/or Nrf2 (Figure 2E). Distinct expression levels of above-selected 16 candidate miRNAs were determined by transcriptome sequencing (Figure 2F), and further validated by real-time qPCR (Figure 2G). On this base in combination of putative Nrf1/2-binding *ARE* motif analysis of those miRNA promoter regions (Figure S9) with a considerable number of their relevant literature [30-35], miR3187-3p and miR1247-5p were hence selected as two key molecules for further study of their specific biological functions.

To verify the biological effects of miR3187-3p and miR1247-5p, *WT* cells were transfected with each of their mimics alongside with a negative control (NC), and then subjected to real-time qPCR analysis of their miRNA expression levels, as well as the mRNA expression abundances of these miRNA-binding downstream target genes (Figure S5), so to explore their inhibitory effects. As shown in Figure 2H (*left panel*), the miR3187-3p expression was significantly upregulated by its mimic to nearly 60 folds than its corresponding control. By contrast, the expression level of miR1247-5p wa salso substantially increased by its mimic so as to reach nearly 6000 folds compared to the NC level (Figure 4I, *left panel*). Interestingly, the miR3187-3p mimic also significantly inhibited MMP19, FN1, SNAI1, TWIST1 and TP53 (all involved in the putative EMT) (Figure 2H, *right panel*). Besides, MMP15 and MMP17 were also substantially inhibited by the miR1247-5p mimic, but with no effects on MMP25 (Figure 2I, *right panel*). From these results, it is inferable that SNAI1, MMP15, MMP17 and MMP25 are much likely to act as key genes involved in the EMT, and thus they were further studied as EMT-related markers in the following experiments. This is also supported by the survival curve of these EMT-marker proteins in clinical data (Figure S4, obtained via http://kmplot.com/analysis/indexwebsite).

### 3.3 Differentially targeting of miR3187-3p and miR1247-5p to SNAI1, MMP15, and MMP17 involved in EMT

To further verify direct roles of miR3187-3p and miR1247-5p in regulating the mRNA expression of *SNAI1, MMP15, MMP17* and *MMP25*, their site-directed mutants on the 3’ -UTRs (Figure 3A) were constructed and subsequently co-transfected with each mimic of miR3187-3p or miR1247-5p into HepG2 cells. The results showed that such miRNA mimic-leading decreases in their cognate target 3’-UTRs-monitored luciferase activity were almost completely blocked by co-transfecting with each corresponding mutants (Figure 3, B to D). However, no significant changes in MMP25-regulated reporter expression were observed, no matter which reporter constructs driven by its wild type or mutant 3’-UTRs had been co-transfected with miR1247-5p (Figure 3E).

**Figure 3.**
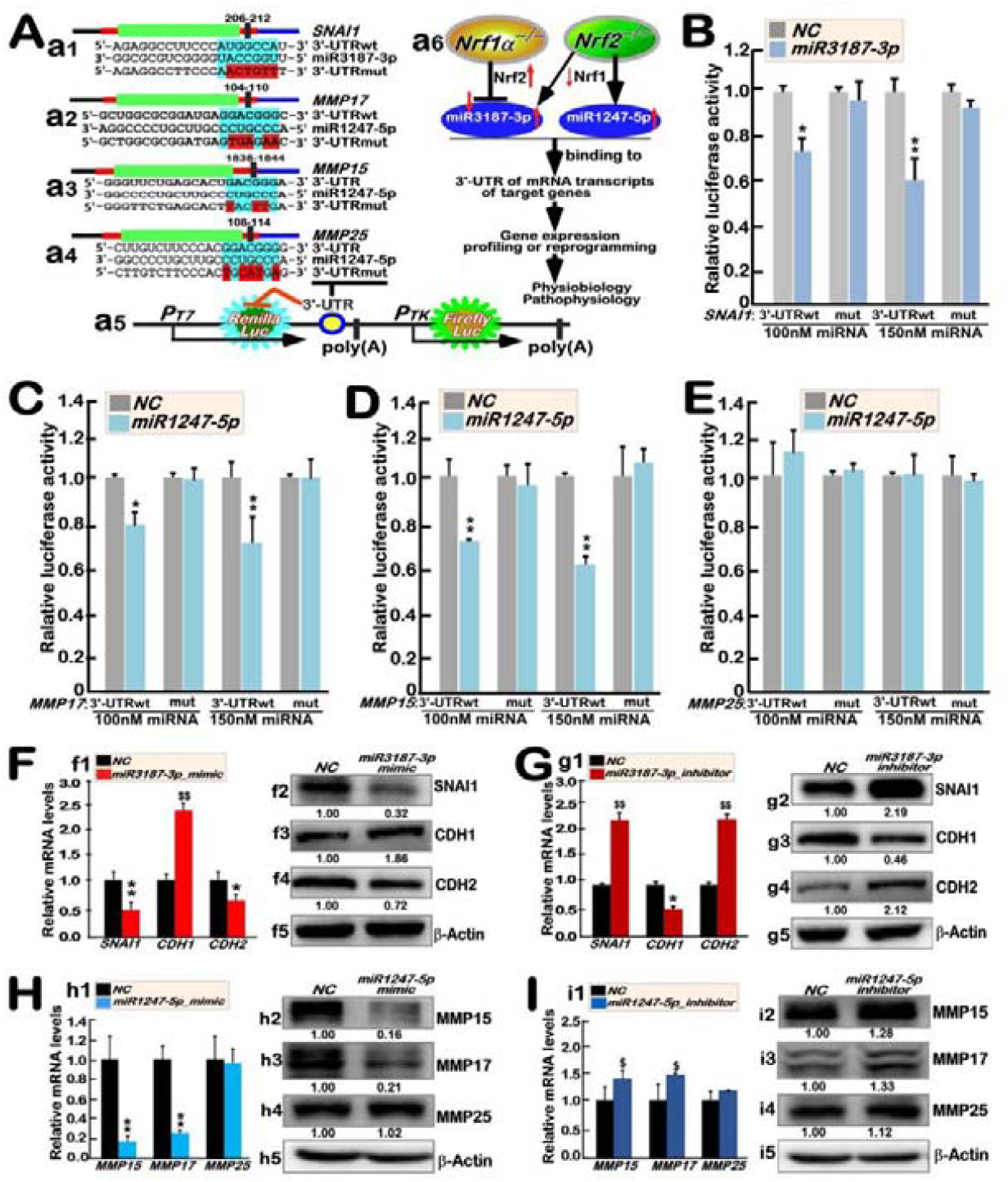
Differential targets of miR3187-3p and miR1247-5p to SNAI1, and MMP15, MMP17, with distinct effects. A. Schematic representation of indicated targets of miR3187-3p and miR1247-5p, together with their mutants, with distinct effects measured by double luciferase reporter constructs used for the following experiments. B to E. HepG2 cells were co-transfected with miR3187-3p, miR1247-5p or their negative controls (NC), along with each of double luciferase reporter constructs entailed by 3′-UTRs of their indicated targets or mutants deciphered above. The 3′-UTRs-modulated luciferase reporter activity regulated by miR3187-3p or miR1247-5p was measured and then subjected to calculation of relative results shown graphically as fold changes (Mean ± S.D.) with significant decreases (*, *p* < 0.05; **, *p* < 0.01), which were determined from three independent experiments each performed in triplicates. F to G. Both mRNA and protein expression levels of indicated target genes regulated by either miR3187-3p or miR1247-5p mimics or inhibitors were determined by real-time qPCR (*left panels*) and Western blotting with indicated primary antibodies (*right panels*). The intensity of all immunoblots was quantified and normalized by that of β-actin, before being shown as fold changes (*on the bottom of indicated blots*). The data of real-time qPCR are also shown graphically as fold changes (Mean ± S.D.) with significant increases ($, *p* < 0.05; $$, *p* < 0.01) and significant decreases (*, *p* < 0.05; **, *p* < 0.01), which were determined from three independent experiments each performed in triplicates.

Next, we further evaluate putative biological effects of miR3187-3p’s and miR1247-5p’s mimics or inhibitors on their downstream EMTrelevant genes by both real-time qPCR and Western blotting, after they had been transfected into two distinct experimental cell lines (HepG2 and Hep3B), respectively. The results showed that the expression of SNAI1 and another mesenchymal marker CDH2 was significantly inhibited by miR3187-3p mimics, but as accompanied by activated expression of the epithelial marker CDH1 at mRNA and protein levels (Figure 3F, *f1 to f4*). By sharp contrast, transfection of miR3187-3p inhibitor (150 nM of its nucleotides) caused a notable increase in the expression of SNAI1 alongside with a promotion in the CDH2 expression, but was also accompanied by significantly inhibited expression of CDH1, the extents of which were inversely correlated with those of elevated SNAI1 (Figure 3G, *g1 to g4*). Furtherly, the miR1247-5p mimics demonstrated a notable inhibitory effect on the expression of MMP15 and MMP17 (Figure 3H, *h1 to h3*), but another increased effect of MMP15 and MMP17 was caused by miR1247-5p inhibitor (Figure 3I, *i1 to i3*). In addition, the MMP25 expression was almost unaffected by miR1247-5p mimic or its inhibitor (Figure 3, H & I). Of note, all the aforementioned data obtained from HepG2 cells were experimentally repeatable in Hep3B cells (as shown in Figure S6).

### 3.4 Differential regulation of targeted EMT genes by distinctive status of miR3187-3p or miR1247-5p

To verify the upstream and downstream effects of SNAI1, we synthesized target-specific siRNA to knockdown SNAI1, along with its downstream CDH1 and CDH2, as shown by both real-time qPCR and Western blotting (Figure 4A). Further examination revealed more significant downregulation of SNAI1 by a miR3187-3p mimic occurring after co-transfection with *siSNAI1* in HepG2 cells, also as accompanied by directly affected expression patterns of its downstream CDH1 and CDH2 (Figure 4B). Conversely, enhanced expression of SNAI1 by miR3187-3p inhibitor was prevented by *siSNAI1* (Figure 4C), which also affected the according expression levels of CDH1 and CDH2. Together, these further corroborated the regulatory relationship between miR3187-3p and SNAI1, and their downstream CDH1 and CDH2.

**Figure 4.**
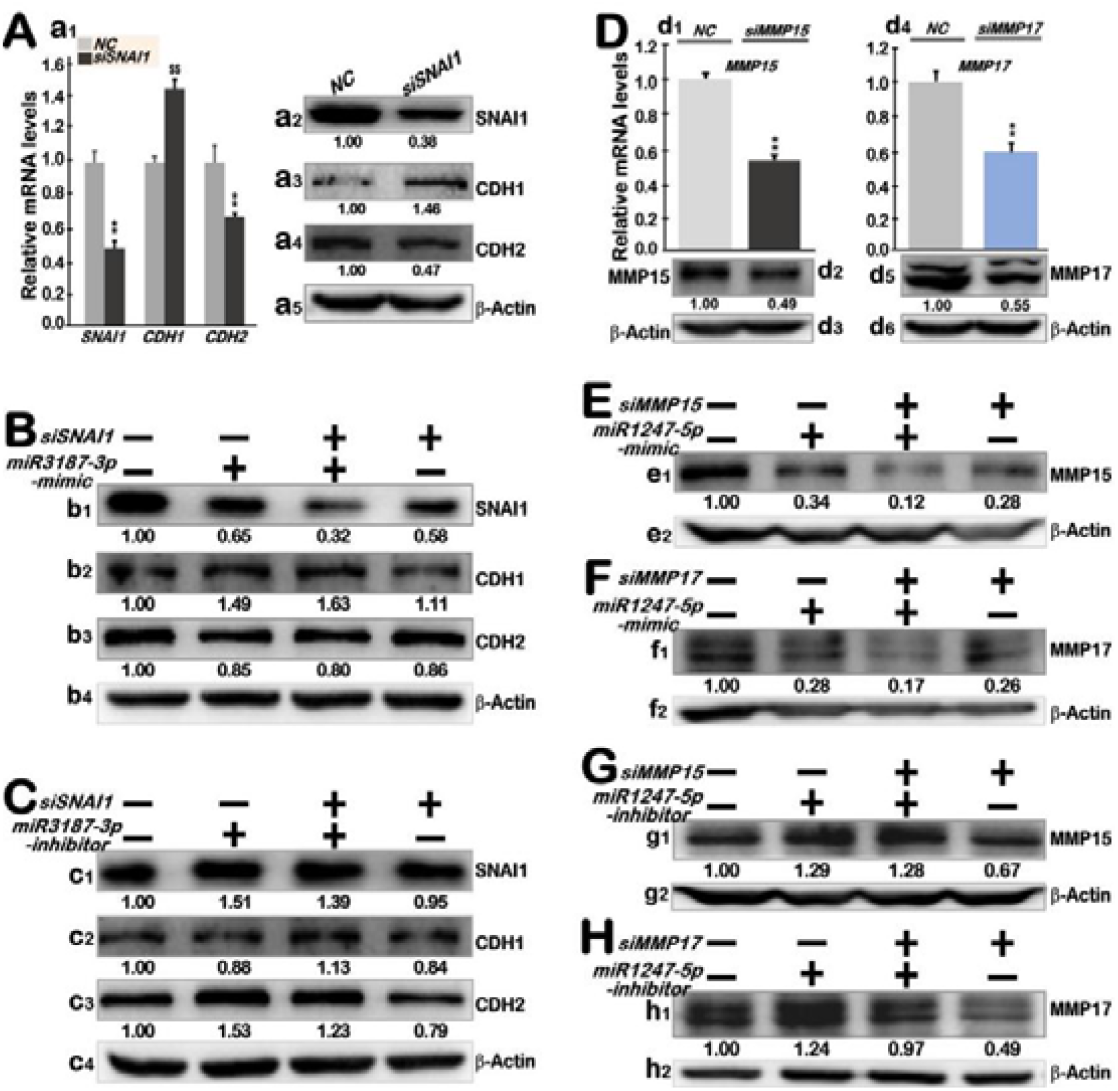
Rescue experiments to verify an upstream or downstream relationship between miRNAs and their targets. A. HepG2 cells that had been transfected with SNAI1-specific siRNA (*siSNAI1*) or negative control (NC) were subjected to real-time qPCR and Western blotting assays to determine both mRNA and protein expression levels of SNAI1 and its downstream CDH1 and CDH2. The qPCR data are shown graphically as fold changes (Mean ± S.D.) with significant increases ($$, *p* < 0.01) and significant decreases (**, *p* < 0.01), which were determined from three independent experiments each performed in triplicates. The intensity of indicated immunoblots was quantified and normalized by that of β-actin, before being shown as fold changes (*on the bottom of indicated blots*). B,C. Two distinct rescue experiments were conducted by co-transfection of siSNAI1 with a miR3187-3p mimic (B) or its inhibitor (C), before the expression abundances of SNAI1 and its downstream CDH1 and CDH2 were determined by immunoblotting with distinct antibodies. The intensity of indicated immunoblots was quantified and normalized by that of β-actin, before being shown as fold changes (*on the bottom of indicated blots*). D. The mRNA and protein expression levels of MMP15 or MMP17 were determined by real-time qPCR Western blotting in HepG2 cells that had been interfered by *siMMP15* or *siMMP17*, respectively. The qPCR data are shown graphically as fold changes (Mean ± S.D.) with significant decreases (**, *p* < 0.01), which were determined from three independent experiments each performed in triplicates. The intensity of indicated immunoblots was quantified and normalized by that of β-actin, before being shown as fold changes (*on the bottom of indicated blots*). E to H. Distinct rescue experiments were carried out by co-transfection of siMMP15 or siMMP17 with a miR1247-5p mimic (E,F) or its inhibitor (G,H), before relevant protein abundances were determined by immunoblotting with distinct antibodies. The intensity of indicated immunoblots was quantified and normalized by that of β-actin, before being shown as fold changes (*on the bottom of indicated blots*).

Next, the downstream regulation of miR1247-5p was verified in a manner similar to the aforementioned approach. That is, the basal expression of MMP15 or MMP17 at their mRNA and protein levels was effectively silenced by *siMMP15* or *siMMP17*, respectively, transfected in HepG2 cells (Figure 4D). Furtherly, markedly reduced expression of MMP15 or MMP17 was observed following co-transfection of *siMMP15* or *siMMP17* with miR1247-5p mimics (Figure 4, E & F), also with a synergistic effect exerted by miR1247-5p mimics with *siMMP15* or *siMMP17*. Conversely, upon co-transfection of miR1247-5p inhibitor with *siMMP15* or *siMMP17*, the inhibitor-leading increases in the expression of MMP15 or MMP17 were partially mitigated by *siMMP15* or *siMMP17*, respectively, hence forming a resistance effect on MMP15 or MMP17 (Figure 4, G &H). Of crucial importance, it is plausible that all the aforementioned data obtained from HepG2 cells (Figure 4) were experimentally repeatable in Hep3B cells (as shown in Figure S7).

In order to further confirm the above-described effects of miR3187-3p or miR1247-5p on their targets, the indicated miRNAs were constructed into the pLKO.1-TRC cloning vector (for their transient expression) or the lentiviral expression system (subjected to their stable expression, along with the negative controls) in HepG2 cells (Figure 5A). As expected, the expression levels of each of indicated miRNAs (i.e., miR3187-3p or miR1247-5p) were substantially incremented to considerably higher extents in all those examined cases, when compared to their negative controls (i.e., NC) (Figure 5, B and C). Further experimental examinations of such highly expressing miRNAs’ effects on their respective targets revealed that both SNAI1 and CDH2 were significantly downregulated by miR3187-3p (no matter whether it had been transiently or stably expressed), as accompanied by enhanced expression of CDH2 (Figure 5, D & F). Similarly, MMP15 and MMP17 were indeed downregulated by miR1247-5p, but as accompanied by unaffected expression of MMP25 (Figure 5, E & G). In addition, all the above-mentioned data were also repeatable in Hep3B cells (as deciphered in Figure S8).

**Figure 5.**
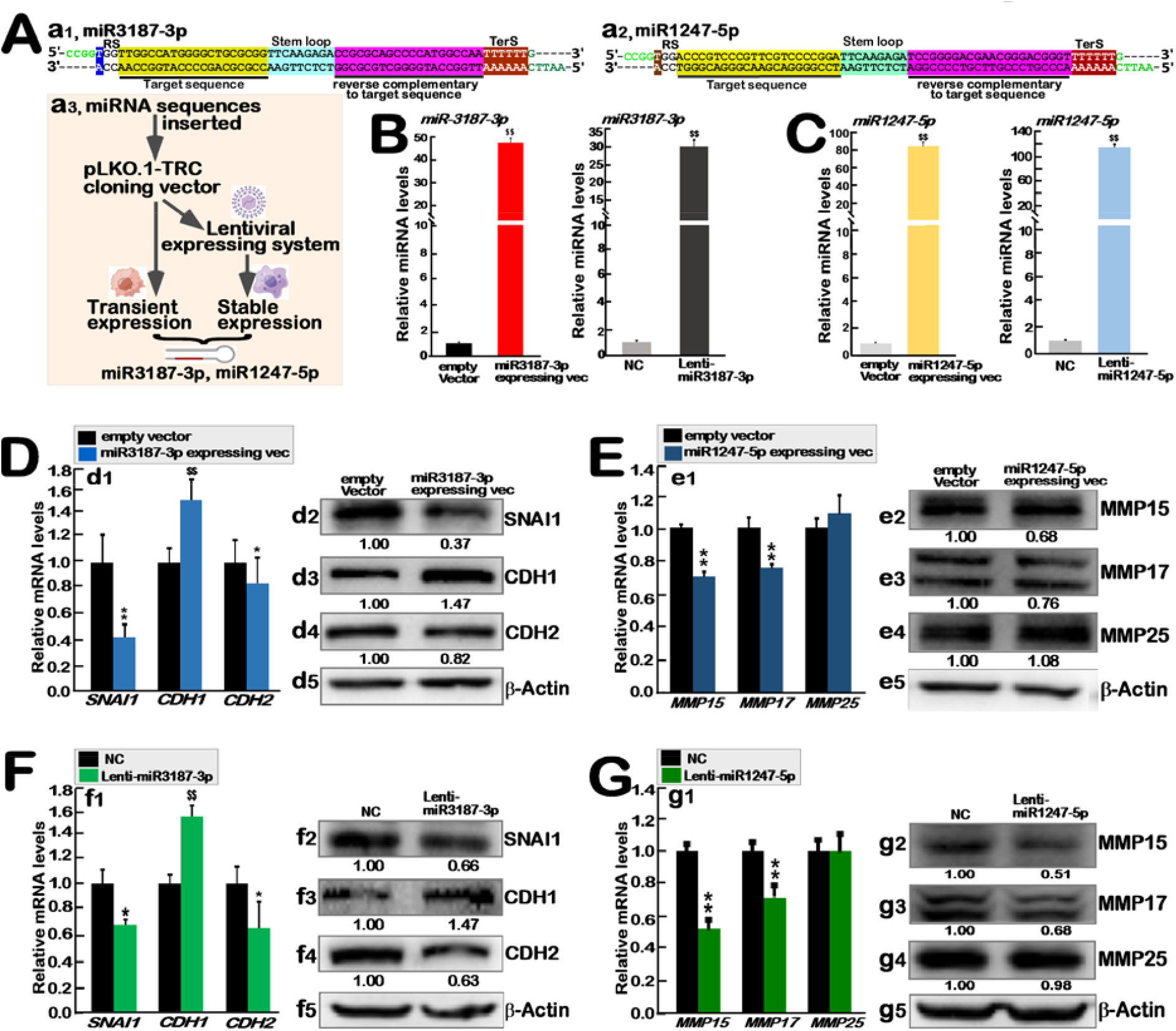
Similar but nuanced effects of transiently or stably expressing miR3187-3p and miR1247-5p on their targets. A. Schematic representations of miR3187-3p or miR1247-5p, that were subjected to transient or stable expression by indicated pLKO.1-TRC vector and Lentiviral expression systems, respectively. B,C. HepG2 cells had been transiently transfected or stably infected with each of the above-indicated miRNAs-expressing systems, before being subjected to real-time qPCR. Their relative expression levels are shown graphically as fold changes (Mean ± S.D.) with significant increases ($$, p < 0.01), which were determined from three independent experiments each performed in triplicates. D,E. The transiently-expressing miR3187-3p’s or miR1247-5p’s effects on their targets were examined by real-time qPCR and Western blotting with indicated antibodies. The qPCR data are shown graphically as fold changes (Mean ± S.D.) with significant increases ($$, *p* < 0.01) and significant decreases (**, *p* < 0.01), which were determined from three independent experiments each performed in triplicates. The intensity of indicated immunoblots was quantified and normalized by that of β-actin, before being shown as fold changes (*on the bottom of indicated blots*). F,G. The stably-expressing miR3187-3p’s or miR1247-5p’s effects on their targets were examined by real-time qPCR and Western blotting with indicated antibodies. The qPCR data are shown graphically as fold changes (Mean ± S.D.) with significant increases ($$, *p* < 0.01) and significant decreases (*, p < 0.05; **, *p* < 0.01), which were determined from three independent experiments each performed in triplicates. The intensity of indicated immunoblots was quantified and normalized by that of β-actin, before being shown as fold changes (*on the bottom of indicated blots*).

### 3.5 Cellular mechanisms by which miR3187-3p and miR1247-5p inhibit the invasive and migratory ability of HCC

Herein, to ascertain the migratory and invasive capabilities of miR3187-3p and miR1247-5p, a series of scratch and transwell experiments were conducted, after miR3187-3p and miR1247-5p had been transiently or stably expressed, in relevant plasmid-transfected or lentivirus-infected HepG2 cells, respectively. As anticipated, the results demonstrated that both miR3187-3p and miR1247-5p are indeed capable of inhibiting the migration and invasion of hepatoma cells to considerably lower extents (Figure 6A, *a1 to a6*). Further investigation of clone proliferation ability of stably-expressing miR3187-3p or miR1247-5p lentivirus-infected cell lines demonstrated that Lenti-miR3187-3p cells exhibited relatively slower growth of hepatoma cells that had been seeded across a range of cell densities (500-1500) (Figure 6C), whilst Lenti-miR-1247-5p cells were only manifested with minimal changes.

**Figure 6.**
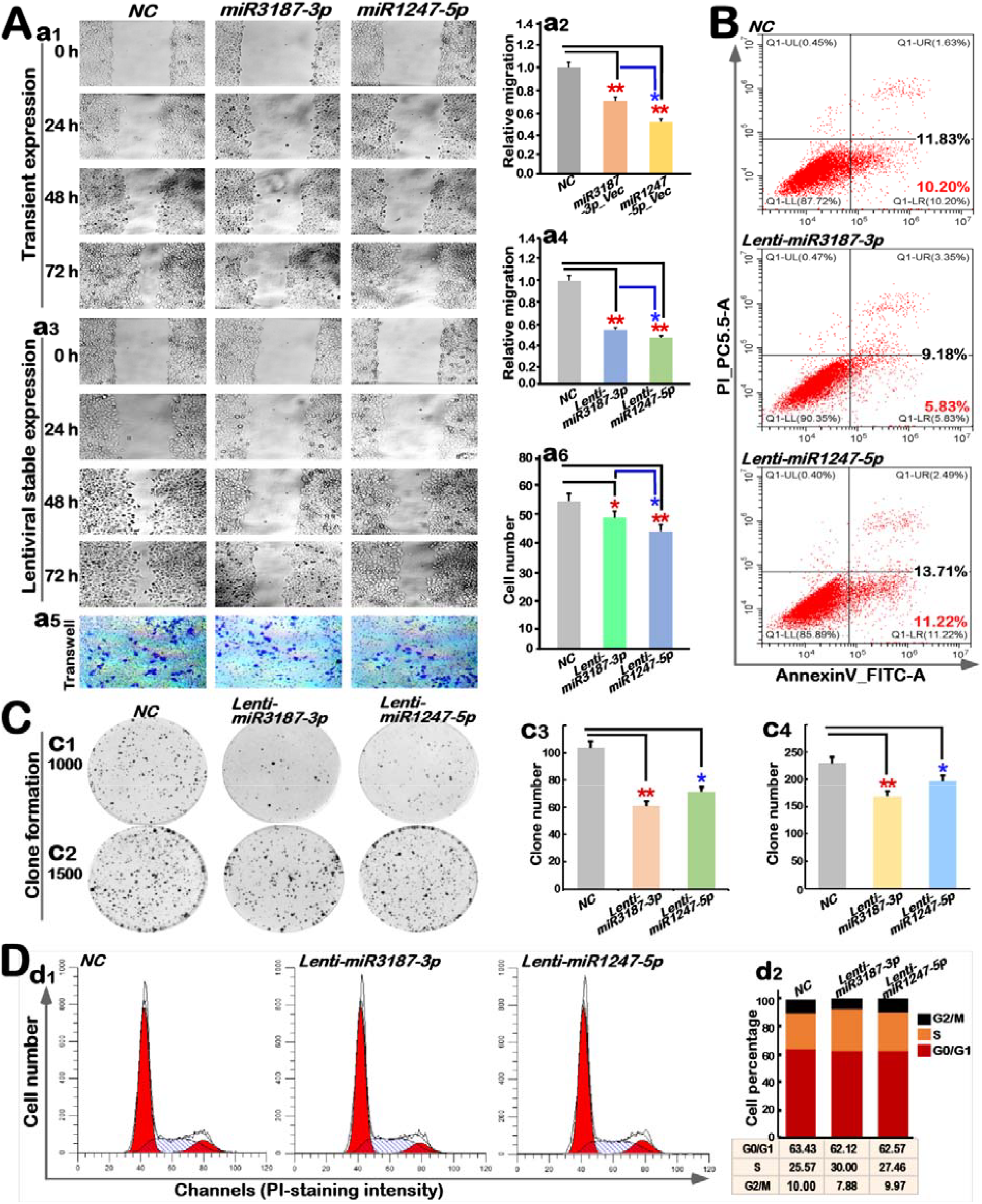
Inhibition of clone proliferation, invasion and migration of hepatoma cells by miR3187-3p and miR1247-5p. A. Scratch and transwell tests revealed that the migration and invasion ability of hepatoma cells were inhibited by miR3187-3p or miR1247-5p, which had been transiently or stably expressed in HepG2 cells. The results were also quantified and shown graphically as fold changes (Mean ± S.D.) with significant decreases (*, p < 0.05; **, *p* <?0.01), which were determined from three independent experiments each performed in triplicates. B. The apoptosis of stably-expressing miR3187-3p or miR1247-5p cell lines was determined by flow cytometry. C. The colony formation assay of stably-expressing miR3187-3p or miR1247-5p cell lines were carried out. The results were quantified and shown graphically as fold changes (Mean ± S.D.) with significant decreases (*, p < 0.05; **, *p* < 0.01), which were determined from three independent experiments each performed in triplicates. D. Flow cytometry analysis of stably-expressing miR3187-3p or miR1247-5p cell line were conducted, as graphically shown by the percentage of distinct cell populations at different phases.

Further insights into apoptosis of stably-expressing miR3187-3p or miR1247-5p hepatoma cells by flow cytometry, the results demonstrated that the former Lenti-miR3187-3p slightly enhanced early apoptosis but reduced late apoptosis (Figure 6B, *middle panel*), whilst the latter Lenti-miR1247-5p can consistently promote early and late apoptosis of HepG2 cells (Figure 6B, *lower panel*), when compare to the controls. Such nuanced effects of miR3187-3p or miR1247-5p on hepatoma cell cyles were further analyzed by flow cytometry. The results unraveled that an increased proportion of cells at the S phase was observed following stable expression of miR3187-3p, as accompanied by another obviously decreased proportion of cells at the G2/M phase, with no changes in the G0/G1 phase proportion (Figure 6D). Rather, stable expression of miR-1247-5p only caused a slight increase in the S phase proportion of hepatoma cells, but also as accompanied by largely unchanged proportions of the cells in the G2/M and G0/G1 phases (Figure 6D), when compared with the controls.

### 3.6 Molecular mechanisms by which distinct miRNAs contribute to the opposing effects of Nrf1 and Nrf2 on EMT

Herein, to ascertain whether Nrf1 exerts a direct influence on miR3187-3p, we constructed a series of luciferase reporter genes driven by 3× *ARE*-adjoining sequences in the miR3187-3p promoter region (as deciphered in Figure 7A). Each of indicated *ARE*-driven reporters was co-transfected into HepG2 cells, together with an expression construct for Nrf1 or empty vector. As expected, the results revealed that Nrf1 exerted notable trans-acting effects on *miR-3187-3p* by its directly transcriptional regulation occurring primarily at its #1 and #4 ARE sites, while additional relative weaker transacting effects were executed by the other two ARE sites of #2 and #5 (Figure 7B).

**Figure 7.**
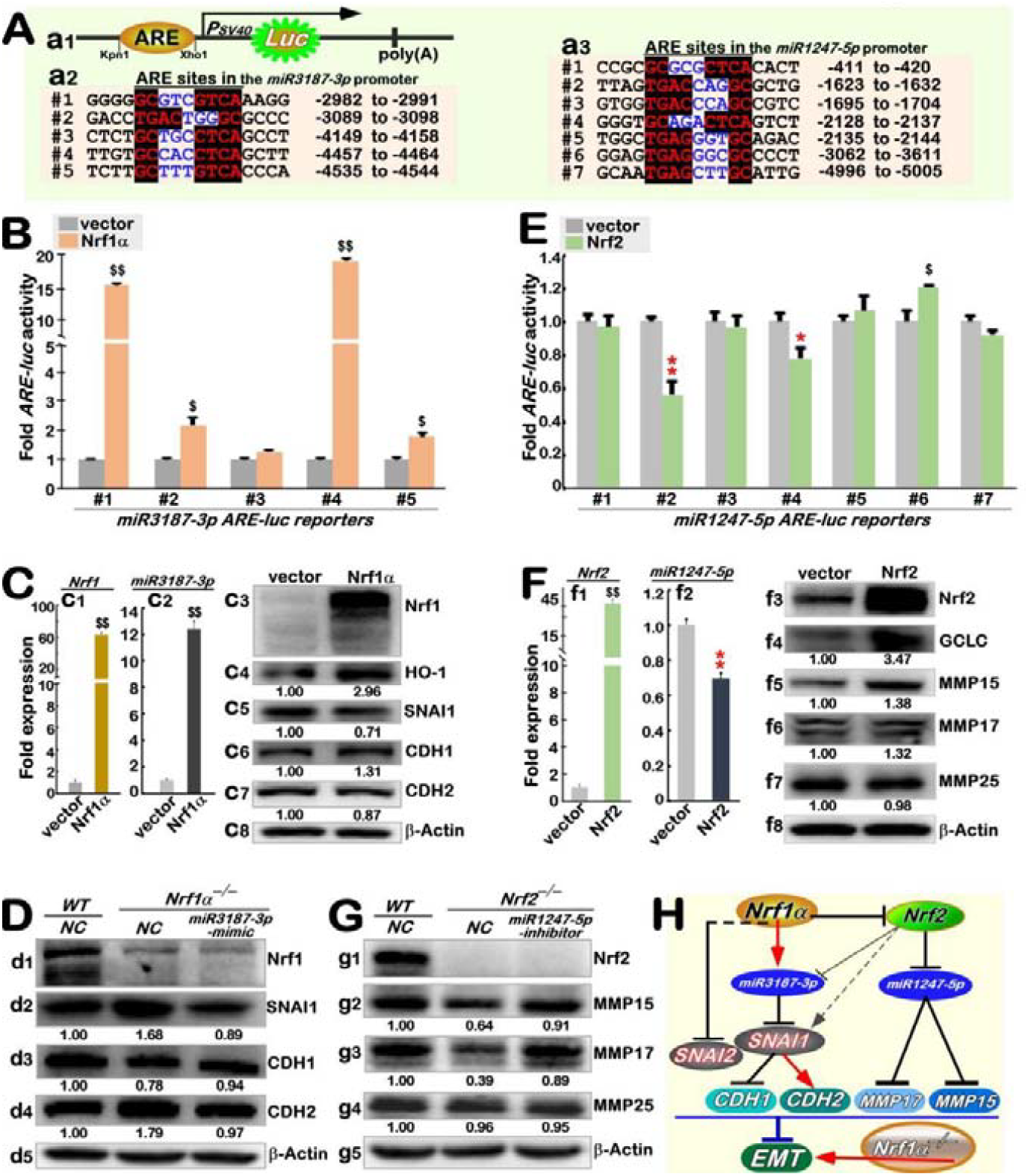
Distinct contributions of miR3187-3p and miR1247-5p to the opposing effects of Nrf1 and Nrf2 on the EMT. B. A schematic representation of ARE-driven luciferase reporters, that were constructed from each of indicated ARE-adjoining sequences within the miR3187-3p and miR1247-5 promoter regions. B. The effects of ectopic Nrf1*α* on the transcriptional activity of miR3187-3p-derived *ARE-luc* reporter genes were determined. The results were graphically shown as fold changes (Mean ± S.D.) with significant increases ($, *p* < 0.05; $$, *p* < 0.01), which were determined from three independent experiments performed in triplicates. C. Distinct effects of ectopic Nrf1*α* on the expression of miR3187-3p and its downstream proteins were determined by real-time qPCR and Western blotting analysis of HepG2 cells allowed for overexpression of Nrf1*α*. The qPCR data are shown graphically as fold changes (Mean ± S.D.) with significant increases ($$, *p* < 0.01), which were determined from three independent experiments each performed in triplicates. The intensity of immunoblots was quantified and normalized by that of β-actin, before being shown as fold changes (*on the bottom of indicated blots*). D. The impact of a miR-3187-3p mimic on the examined proteins in *Nrf1α*^*−/−*^ cells was determined by Western blotting of both *WT* and *Nrf1α*-deficient cell lines that had been transfected with this miRNA mimic or its negative control (NC). The intensity of immunoblots was quantified and normalized by that of β-actin, before being shown as fold changes (*on the bottom of indicated blots*). E. Distinct effects of ectopic Nrf2 on the transcriptional activity of miR1247-5p-derived ARE-luc reporter genes were determined. The resulting data were graphically shown as fold changes (Mean ± S.D.) with a significant increase ($$, *p* < 0.01) and another significant decrease (*, *p* < 0.05; **, *p* < 0.01), each of which were determined from three independent experiments performed in triplicates. F. Distinct effects of ectopic Nrf2 on the expression of miR1247-5p and its downstream proteins were determined by real-time qPCR and Western blotting analysis of HepG2 cells allowed for overexpression of Nrf2. The qPCR data are shown graphically as fold changes (Mean ± S.D.) with a significant increase ($$, *p* < 0.01) and another significant decrease (**, *p* < 0.01), which were determined from three independent experiments each performed in triplicates. The intensity of immunoblots was quantified and normalized by that of β-actin, before being shown as fold changes (*on the bottom of indicated blots*). G. The impact of a miR1247-5p inhibitor on the examined proteins in *Nrf2*^*−/−*^ cells was determined by Western blotting of both *WT* and *Nrf2*-deficient cell lines that had been transfected with this miRNA inhibitor or its negative control (NC). The intensity of immunoblots was quantified and normalized by that of β-actin, before being shown as fold changes (*on the bottom of indicated blots*). H. A model is proposed to give a concise explanation of the finding that the EMT of HCCs is negatively regulated by Nrf1*α*-activating miR3187-3p, but also positively promoted by Nrf2-inhibiting miR1247-5p. For detailed descriptions, please see supplemental Figure S11.

Next, we further examined effects of Nrf1-forced expression on miR3187-3p and its downstream targets. The results showed a significant increase in the miR3187-3p expression caused by over-expression of Nrf1*α* in HepG2 cells (Figure 7C, *c2 vs c1*). Of particular interest, it was observed that overexpression of Nrf1*α* led to the evident inhibition of SNAI1 and CDH2 (Figure 7C, *c3*), but also was accompanied by substantial activation of both CDH1 and HO-1 (as a canonical downstream enzyme of Nrf1 to exert its antioxidant and detoxifying responses). Further examinations of miR3187-3p’s effects on the EMT-relevant genes in *Nrf1α*^*−/−*^ cells revealed that this miR3187-3p mimic was capable of attenuating the elevation of SNAI1 caused by loss of *Nrf1α*, so that reduced SNAI1 also led to the restoration of CDH1 abundances, along with another reduction in CDH2 expression, when compared to their relevant controls (Figure 7D).

Similar experiments were performed as described above, to verify a direct effect of Nrf2 on miR1247-5p. A series of *ARE*-driven luciferase reporter genes were constructed from the *miR1247-5p* promoter region (Figure 7A, *a3*). The results demonstrated that Nrf2 exerted a pronounced inhibitory effect on miR1247-5p at its two ARE sites of #2 and #4 within its promoter region (Figure 7E), but as accompanied by a modest transacting effect on its #6 ARE site. The latter weak effect appeared to be nearly inconsequential, due to a quite far distance of the 6^th^ ARE from its transcription start site. This is supported by further experiments showing that overexpression of Nrf2 led to an overall significant inhibitory effect on the miR-1247-5p expression in HepG2 cells (Figure 7F, *cf. f2 with f1*). Such reduction of miR-1247-5p by ectopic Nrf2 caused obvious increases in the expression of MMP15 and MMP17, but not MMP25, aside from transactivation of the cognate downstream target GCLC by this CNC-bZIP factor (Figure 7F, *f4 to f6*). Conversely, following the transfection of a miR1247-5p inhibitor into *Nrf2* ^*−/−*^ cells, it was shown that this miRNA inhibitor enabled to mostly rescue the declined expression of MMP15 and MMP17 caused by loss of *Nrf2* (Figure 7G), but no any effects were observed on MMP25. Of note, the above-mentioned experimental data were all repeatable to be obtained from Hep3B cells (as shown in Figure S10). Taken together, these demonstrated that the EMT of HCCs is negatively regulated by Nrf1*α*-activating miR3187-3p, but positively promoted by Nrf2-inhibiting miR1247-5p (Figures 7H and S11).

## 4. Discussion

Human liver cancer is one of the most prevalent malignant neoplasms worldwide with a high incidence of morbidity and mortality. Of pivotal significance, it is how to overcome the metastasis of hepatocellular carcinoma (HCC), because it represents a crucial step in advancing the treatment of this disease. For this end, it is of importance to elucidate the cellular and molecular mechanisms whereby the EMT exerts a critical role in the promotion of HCC initiation, progressive development and malignant metastasis. For instance, by suppression of the E-cadherin (CDH1) expression during EMT, it can indeed enable for transformation of a typical polygonal, cobblestone-like epithelial cell shape to acquire another spindle-type mesenchymal morphology. As a consequence, this is also allowed to acquire the additional ability to invade and metastasize remotely. In this process, a progressive increased expression pattern of those mesenchymal-associated markers (including N-calmodulin, vimentin and fibronectin, as well as matrix metallopeptidease proteins (MMPs)[29]) is accompanied by another decreased expression pattern of the epithelial-associated proteins. Overall, these prior studies have demonstrated that a canonical EMT has been successfully achieved through relevant transcription factors, including Snail, Twist, and ZEB families, which exert a dominant regulatory influence to tightly govern the expression abundances of both epithelial and mesenchymal markers. In this study, it is found that the EMT of HCCs is negatively regulated by antioxidant transcription factor Nrf1*α*-activating miR3187-3p signaling, but also positively promoted by its homological factor Nrf2-inhibiting miR1247-5p network (as illustrated in Figure 7H, and also see Figure S11).

Since the EMT is dictated by the expression profiling of specific genes regulated by relevant transcription factors, it is inferable that microRNAs also play a pivotal role in the transcriptional regulation of those indispensable genes for EMT[36, 37]. A previous study demonstrated that the forced expression of miR194 in hepatic stromal cells results in an reduction of N-calmodulin, which, in turn, prevents the cell migration and invasion [12]. Furthermore, miR148a has also been shown to reduce the accumulation of SNAI1 by binding to Met (a proto-oncogene-type receptor tyrosine kinase) in hepatocyte growth, which enables to activate the downstream Akt phosphorylation at its Ser^473^ but also inhibit the phosphorylation of GSK-3β at Ser^9^, thereby leading to remission of EMT in HCCs [38]. Besides, it has also been proposed that Nrf2 plays a role in the EMT process of liver cancer cells [27, 39], although antioxidant transcription factors Nrf1 and Nrf2 are known to play a vital role in maintaining redox homeostasis and organ integrity across within organisms [25, 40]. Nevertheless, the involvement of Nrf1- and Nrf2-mediated microRNAs in EMT regulation in liver cancer cells remains to be not yet well documented, as far as we know from the current literature. In this study, we investigated the differential roles of miRNAs regulated by Nrf1 and Nrf2 in putative EMT of HCC cells, along with such a novel view to elucidate the impact of these regulatory processes on the EMT of liver cancer cells. The cellular-molecular mechanisms accounting for the phenotypic differences between *Nrf1α*^*−/−*^ and *Nrf2*^*−/−*^ cell lines derived from their *WT* cells were further verified by bioinformatics analysis of small RNA-sequencing and transcriptome, in combination with the classic ‘wet’ experiments.

Here is a major object of this study to aim at ascertain how miR3187-3p and miR1247-5p are regulated by Nrf1 and/or Nrf2, respectively. Amongst those accurately screened downstream targets of miR3187-3p and miR1247-5p, SNAI1, MM15, MMP17, and MMP25 were determined as the key EMT molecules for further investigation. Subsequently, the functional effects of miR3187-3p and miR1247-5p on EMT were further identified by administering their mimics and inhibitors. The mechanisms underlying the action of miR3187-3p and miR1247-5p were explored in-depth by using their transient or stable expression cell lines. As expected, the results demonstrated that miR3187-3p could target and inhibit the expression of SNAI1, as described elsewhere [9] and that, as a typical EMT-related transcription factor, SNAI1 could precision-regulate the expression of its downstream CDH1 and CDH2 to certain homeostatically balanced extents, thereby inhibiting EMT. By contrast, miR1247-5p has been shown to directly target both MM15 and MP17, reducing the invasive capacity of liver cancer cells and hence inhibiting EMT. Such differential regulation of miR3187-3p and miR1247-5p by Nrf1 and/or Nrf2 and their distinct contributions to the opposing effects of these two CNC-bZIP factor on the EMT of liver cancer cells. Altogether, the evidence has been provided herein, revealing that miR3187-3p is positively regulated by Nrf1, whilst miR1247-5p is negatively regulated by Nrf2. Therefore, it is plausible that the occurrence of EMT in liver cancer cells is tightly confined by Nrf1-activating miR3187-3p towards the SNAI1-CDH1/2 signaling axis, but promoted by Nrf2-inhibiting miR1247-5p signaling to the MMP15/17 network.

Such two key miRNAs, miR3187-3p and miR1247-5p identified in this study, could be characterized to function as tumour suppressors, which is rather not in contradiction with previous reports [41-43]. This is also further supported by prior researches demonstrating that the miR3187-3p expression is significantly diminished in many malignant tumours, while the elevated levels of miR1247-5p are linked to those favorable patient prognoses. Thereby, it is inferable that Nrf1, which also acts as a putative tumour suppressor, can regulate the EMT process of HCC cells by modulating miR3187-3p in addition to controlling robust redox homeostasis. By sharp contrast, Nrf2, as a crucial redox regulator within cells, has been demonstrated by numerous studies and clinical data analysis to exhibit aberrant expression of this CNC-bZIP factor, resulting in malignant proliferation of many tumours, with a poor prognosis [27]. Herein, Nrf2 has been shown to inhibit the expression of miR1247-5p, which, in turn, can promote the EMT process of liver cancer. Taken together, both miR3187-3p and miR1247-5p enable to function as intermediate bridges of Nrf1 and/or Nrf2 between their regulatory networks involved in the activation of EMT signaling towards HCC progression (Figure S11).

As such being the case, a limitation of this study should also be declared, that all the relevant researches have been conducted primarily at the cellular and molecular levels. Therefore, it is imperative for us to develop animal translational models, aiming to elucidate the consequences of animal experimentation in light of the discrepancies between *in vivo* and *in vitro* experiments. This will be a key focus of our future research to explore distinct contributions of miR3187-3p and miR1247-5p to the opposing effects of Nrf1 and Nrf2 on the EMT during cancer development and malignance.

## Supporting information

sFigureS1

Table S1 Primers for qPCR

Table S2 Primers for plasmid construction

Table S3 Information on the antibodies

Table S4 The cell lines utilized in this study

Table S5 Glossary of abbreviations

## Ethics approval and consent to participate

Not applicable.

## Consent for publication

Not applicable.

## Availability of data and materials

All data needed to evaluate the conclusions in the paper are present in this publication along with the supplementary documents that can be found online. Additional other data related to this paper may also be requested from the corresponding author (with a lead contact at the, or).

## Author contributions

J.C., J.F., Y-p.Z., and S.H. performed all the experiments with help of M.W., collected all the relevant data, and made the manuscript draft with figures and supplemental information. M.W. did bioinformatics analysis of datasets. Lastly, Y.Z. designed and supervised this study, analyzed all the data, helped to prepare all figures with cartoons, wrote and revised the paper.

## Acknowledgments

We thank to all those present and past members of Prof. Zhang’s laboratory (at Chongqing University, China) for giving critical discussion and invaluable help with this work.

## References

1. Sung, H., et al., Global Cancer Statistics 2020: GLOBOCAN Estimates of Incidence and Mortality Worldwide for 36 Cancers in 185 Countries. CA Cancer J Clin, 2021. 71(3): p. 209–249.

2. Acloque, H., J.P. Thiery, and M.A. Nieto, The physiology and pathology of the EMT. Meeting on the epithelial-mesenchymal transition. EMBO Rep, 2008. 9(4): p. 322–6.

3. Trelstad RLH.E., Revel JD, Cell contact during early morphogenesis in the chick embryo. Developmental Biology, 1967. 16(1): p. 78–106.

4. Kong, W., et al., MicroRNA-155 is regulated by the transforming growth factor beta/Smad pathway and contributes to epithelial cell plasticity by targeting RhoA. Mol Cell Biol, 2008. 28(22): p. 6773–84.

5. Su, H., et al., MicroRNA301a targets WNT1 to suppress cell proliferation and migration and enhance radiosensitivity in esophageal cancer cells. Oncol Rep, 2019. 41(1): p. 599–607.

6. Ma, L., et al., miR-9, a MYC/MYCN-activated microRNA, regulates E-cadherin and cancer metastasis. Nat Cell Biol, 2010. 12(3): p. 247–56.

7. Qi, J., et al., MiR-365 regulates lung cancer and developmental gene thyroid transcription factor 1. Cell Cycle, 2012. 11(1): p. 177–86.

8. Lv, Z.D., et al., miR-655 suppresses epithelial-to-mesenchymal transition by targeting Prrx1 in triple-negative breast cancer. J Cell Mol Med, 2016. 20(5): p. 864–73.

9. Lamouille, S., J. Xu, and R. Derynck, Molecular mechanisms of epithelial-mesenchymal transition. Nat Rev Mol Cell Biol, 2014. 15(3): p. 178–96.

10. Kim, T.W., et al., MicroRNA-17-5p regulates EMT by targeting vimentin in colorectal cancer. British Journal of Cancer, 2020. 123(7): p. 1123–1130.

11. Zhang, L., et al., MicroRNA-10b Triggers the Epithelial-Mesenchymal Transition (EMT) of Laryngeal Carcinoma Hep-2 Cells by Directly Targeting the E-cadherin. Appl Biochem Biotechnol, 2015. 176(1): p. 33–44.

12. Meng, Z., et al., miR-194 is a marker of hepatic epithelial cells and suppresses metastasis of liver cancer cells in mice. Hepatology, 2010. 52(6): p. 2148–57.

13. Dongre, A. and R.A. Weinberg, New insights into the mechanisms of epithelial-mesenchymal transition and implications for cancer. Nat Rev Mol Cell Biol, 2019. 20(2): p. 69–84.

14. Viola Vargová, M.P., and Viola Mechírová, Matrix Metalloproteinases. 2012: Current Biology. p. 692.

15. Li, J., et al., MicroRNA-19 triggers epithelial-mesenchymal transition of lung cancer cells accompanied by growth inhibition. Lab Invest, 2015. 95(9): p. 1056–70.

16. Zhang, L., Y. Liao, and L. Tang, MicroRNA-34 family: a potential tumor suppressor and therapeutic candidate in cancer. J Exp Clin Cancer Res, 2019. 38(1): p. 53.

17. Hornstein, E. and N. Shomron, Canalization of development by microRNAs. Nat Genet, 2006. 38 Suppl: p. S20–4.

18. Ru, P., et al., miRNA-29b suppresses prostate cancer metastasis by regulating epithelial-mesenchymal transition signaling. Mol Cancer Ther, 2012. 11(5): p. 1166–73.

19. anru Zhao, Z.Y., Huaping Li,Jiahui Fan,Shenglan Yang, Chen Chen and Dao Wen Wang, MiR-30c protects diabetic nephropathy by suppressing__epithelial-to-mesenchymal transition in db_db mice. Aging Cell, 2017. 16(2): p. 387–400.

20. Dong, P., et al., MiR-137 and miR-34a directly target Snail and inhibit EMT, invasion and sphere-forming ability of ovarian cancer cells. J Exp Clin Cancer Res, 2016. 35(1): p. 132.

21. Fazilaty, H., et al., A gene regulatory network to control EMT programs in development and disease. Nat Commun, 2019. 10(1): p. 5115.

22. Siemens, H., et al., miR-34 and SNAIL form a double-negative feedback loop to regulate epithelial-mesenchymal transitions. Cell Cycle, 2011. 10(24): p. 4256–71.

23. Wang, T., et al., miR-15a-3p and miR-16-1-3p Negatively Regulate Twist1 to Repress Gastric Cancer Cell Invasion and Metastasis. Int J Biol Sci, 2017. 13(1): p. 122–134.

24. Rupaimoole, R. and F.J. Slack, MicroRNA therapeutics: towards a new era for the management of cancer and other diseases. Nat Rev Drug Discov, 2017. 16(3): p. 203–222.

25. Zhang, Y. and Y. Xiang, Molecular and cellular basis for the unique functioning of Nrf1, an indispensable transcription factor for maintaining cell homoeostasis and organ integrity. Biochem J, 2016. 473(8): p. 961–1000.

26. Meng Wang, Y.R., Shaofan Hu, Keli Liu, Lu Qiu and Yiguo Zhang TCF11 Has a Potent Tumor Repressing Effect Than Its Prototypic Nrf1a by Definition of Both Similar Yet Different Regulatory Profiles, With a Striking Disparity From Nrf2. 2021 Front Oncol: Front Oncol. p. 707032.

27. Kai Yazaki, Y.M., Kazufumi Yoshida, Mingma Sherpa, Masayuki Nakajima, Masashi Matsuyama, Takumi Kiwamoto, Yuko Morishima, Yukio Ishii, Nobuyuki Hizawa, ROS-Nrf2 pathway mediates the development of TGF-β1-induced epithelial-mesenchymal transition through the activation of Notch signaling. 2021, European Journal of Cell Biology. p. 151181.

28. Brenda J. Reinhar Frank J. Slack, M.B., Amy E. Pasquinelli, Jill C. Bettinger, Ann E. Rougvie, and H.R.H.G. Ruvkun, The 21-nucleotide let-7 RNA regulates developmental timing in Caenorhabditis elegans. nature, 2000. 403: p. 901–906.

29. Nieto, M.A., et al., Emt: 2016. Cell, 2016. 166(1): p. 21–45.

30. Ashrafizadeh, M., et al., MicroRNAs and Their Influence on the ZEB Family: Mechanistic Aspects and Therapeutic Applications in Cancer Therapy. Biomolecules, 2020. 10(7): p. 1040.

31. Kim, T., et al., p53 regulates epithelial-mesenchymal transition through microRNAs targeting ZEB1 and ZEB2. J Exp Med, 2011. 208(5): p. 875–83.

32. Zheng, T., et al., Functional mechanism of hsa-miR-128-3p in epithelial-mesenchymal transition of pancreatic cancer cells via ZEB1 regulation. PeerJ, 2022. 10: p. e12802.

33. Zhang, X., et al., miR-498 inhibits the growth and metastasis of liver cancer by targeting ZEB2. Oncol Rep, 2019. 41(3): p. 1638–1648.

34. Yin, L.C., et al., MicroRNA-361-5p Inhibits Tumorigenesis and the EMT of HCC by Targeting Twist1. Biomed Res Int, 2020. 2020: p. 8891876.

35. Lu, J., et al., MiR-205 suppresses tumor growth, invasion, and epithelial-mesenchymal transition by targeting SEMA4C in hepatocellular carcinoma. FASEB J, 2018: p. fj201800113R.

36. Feng, J., et al., The Role of MicroRNA in the Regulation of Tumor Epithelial-Mesenchymal Transition. Cells, 2022. 11(13).

37. Garinet, S., et al., Clinical assessment of the miR-34, miR-200, ZEB1 and SNAIL EMT regulation hub underlines the differential prognostic value of EMT miRs to drive mesenchymal transition and prognosis in resected NSCLC. British Journal of Cancer, 2021. 125(11): p. 1544–1551.

38. Liu, M., et al., Hypoxia-induced activation of Twist/miR-214/E-cadherin axis promotes renal tubular epithelial cell mesenchymal transition and renal fibrosis. Biochem Biophys Res Commun, 2018. 495(3): p. 2324–2330.

39. Petrelli, A., et al., MicroRNA:gene profiling unveils early molecular changes and nuclear factor erythroid related factor 2 (NRF2) activation in a rat model recapitulating human hepatocellular carcinoma (HCC). Hepatology, 2014. 59(1): p. 228–41.

40. Younossi, Z., et al., Global burden of NAFLD and NASH: trends, predictions, risk factors and prevention. Nature Reviews Gastroenterology & Hepatology, 2017. 15(1): p. 11–20.

41. Vignard, V., et al., MicroRNAs in Tumor Exosomes Drive Immune Escape in Melanoma. Cancer Immunology Research, 2020. 8(2): p. 255–267.

42. Huang, C., et al., PTBP1-mediated biogenesis of circATIC promotes progression and cisplatin resistance of bladder cancer. International Journal of Biological Sciences, 2024. 20(9): p. 3570–3589.

43. Mauro Scaravilli, K.P.P., Anniina Brofeldt. MiR-1247-5p is overexpressed in castration resistant prostate cancer and targets MYCBP2. Prostate, 2015. 8(75): p. 8.

