## Supplementary figures and images for "The opposing mechanisms by which miRNAs critically contribute to differential roles of Nrf1 and Nrf2 in modulating the epithelial-mesenchymal transformation of hepatocellular carcinoma"

### FigureS2

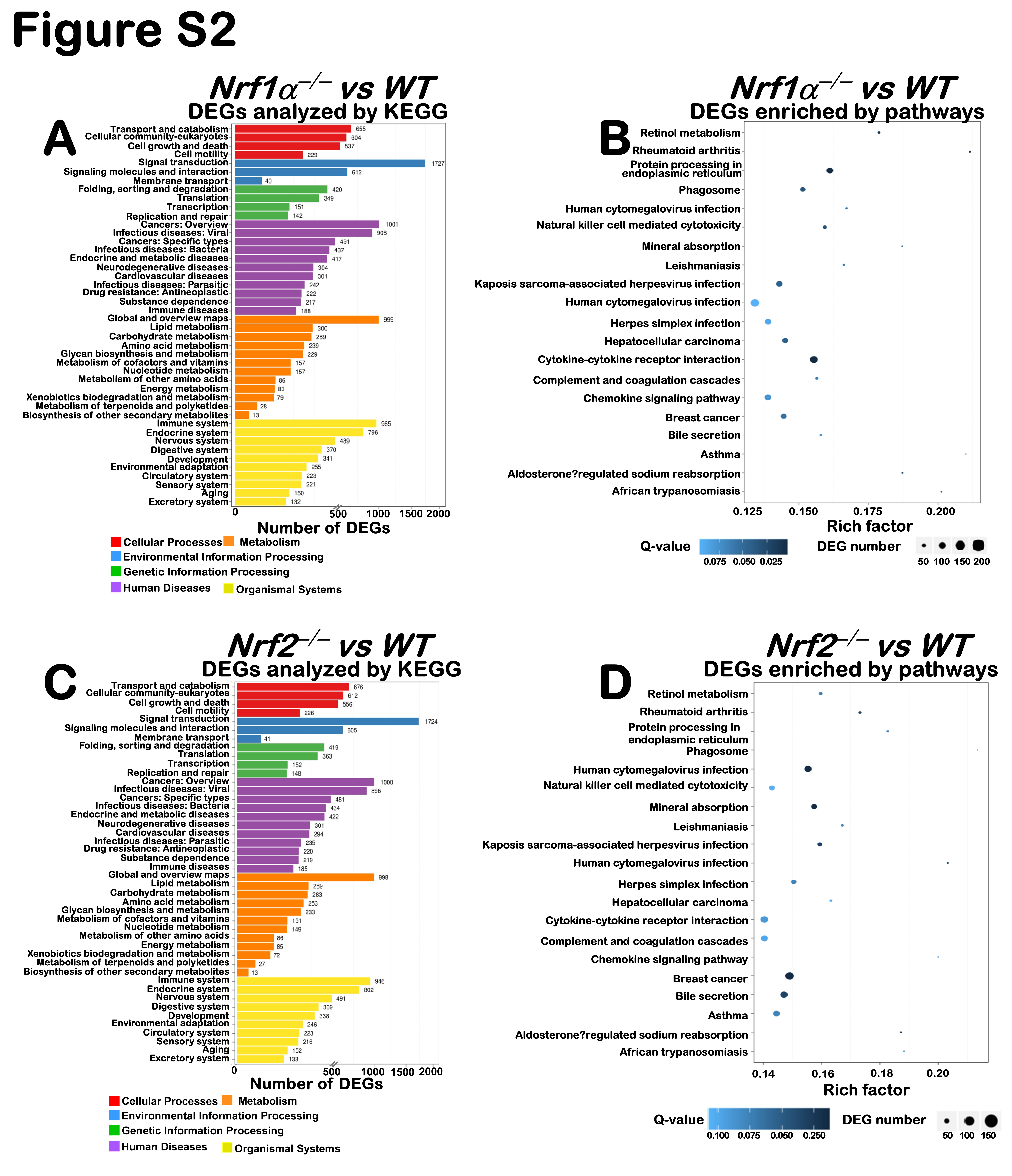

### FigureS3

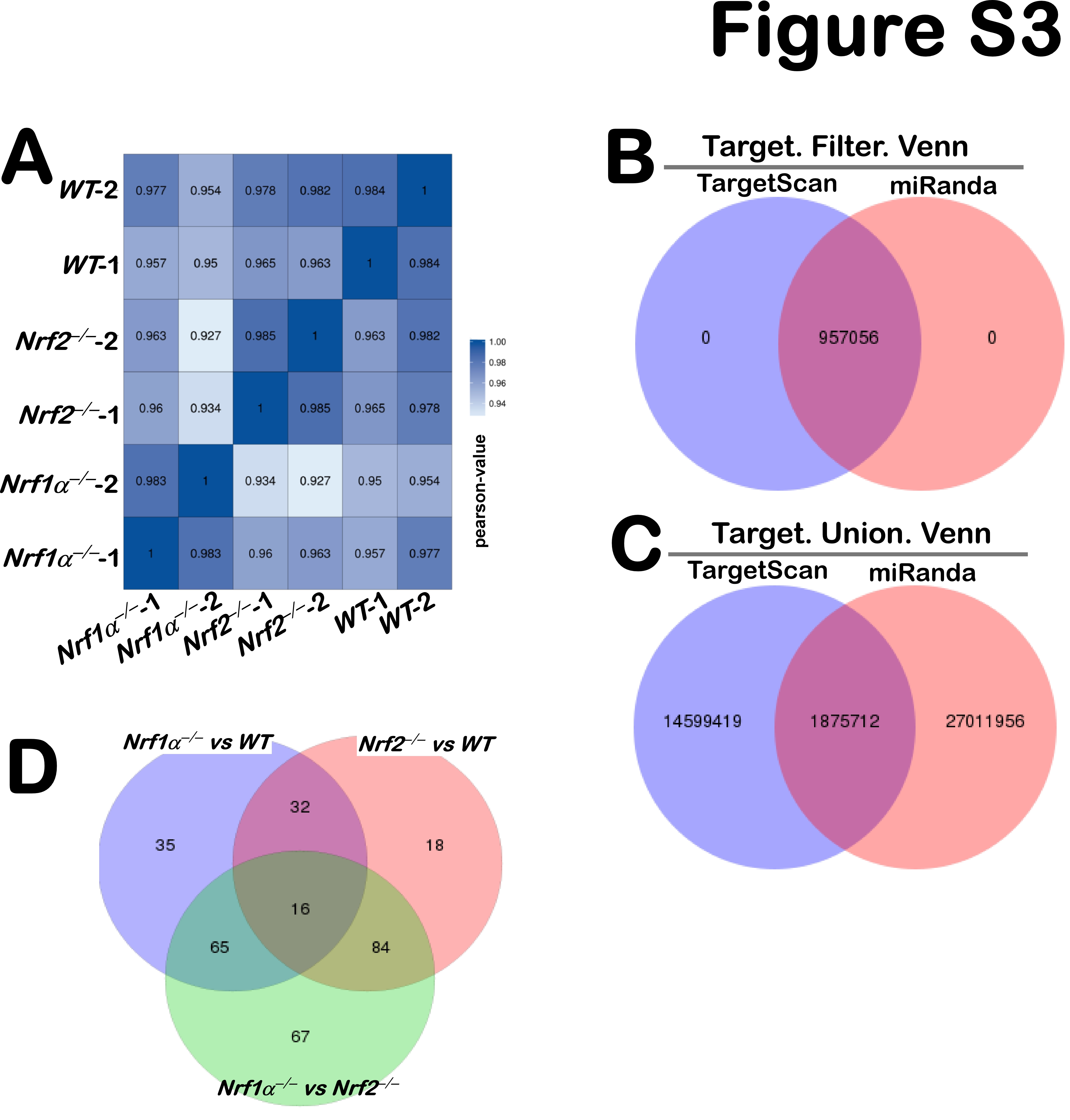

### FigureS4

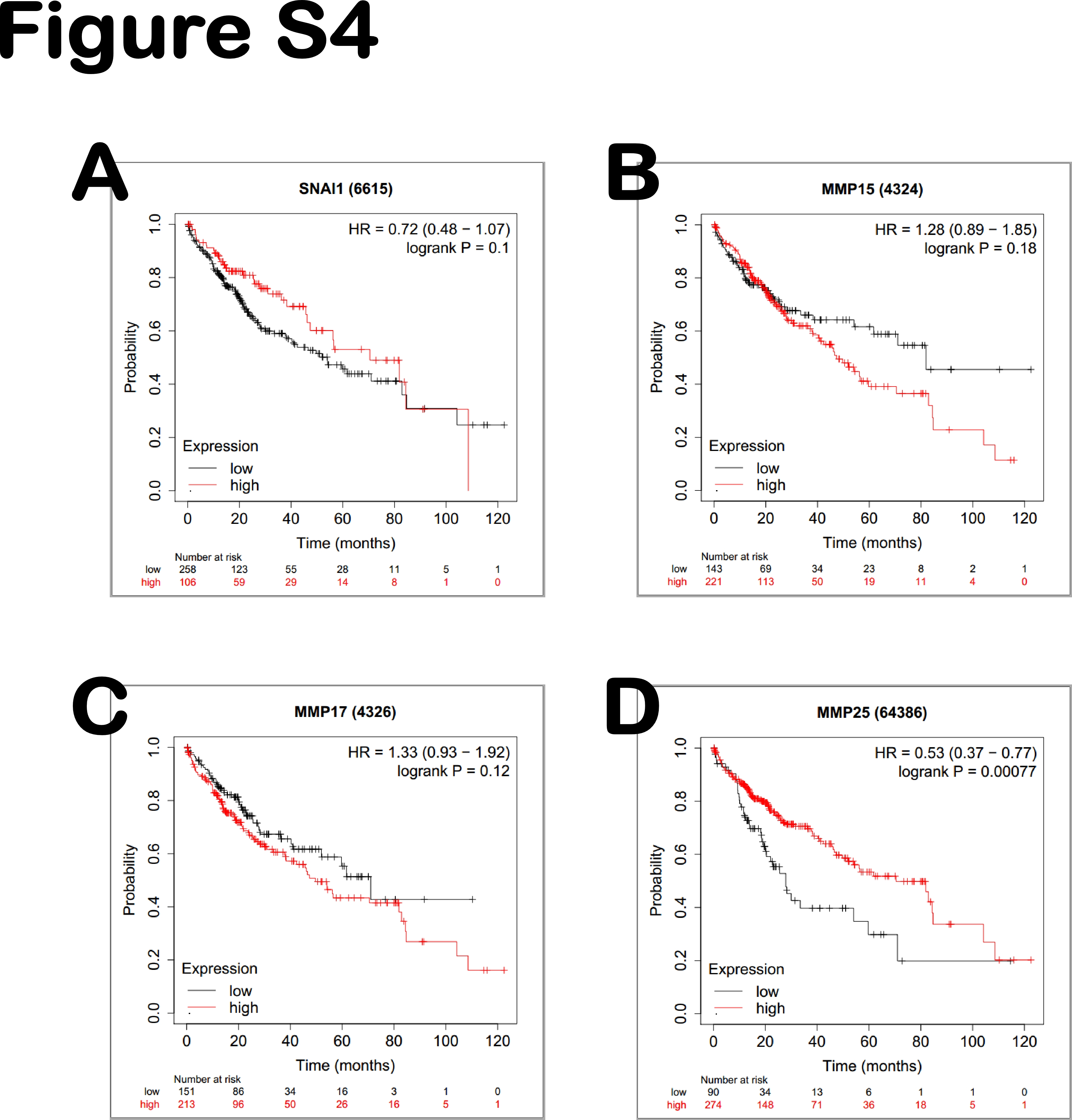

### FigureS5

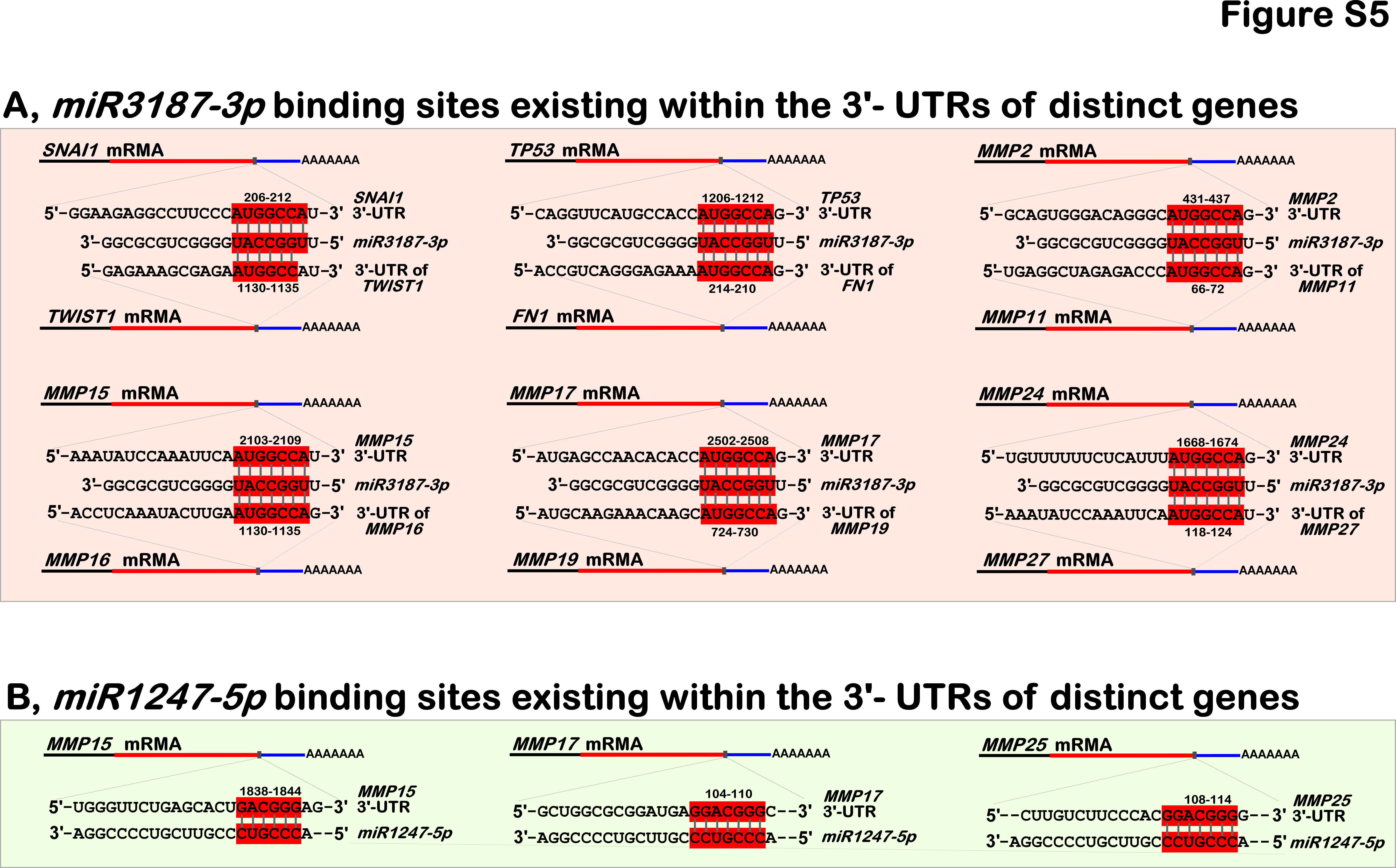

### FigureS6

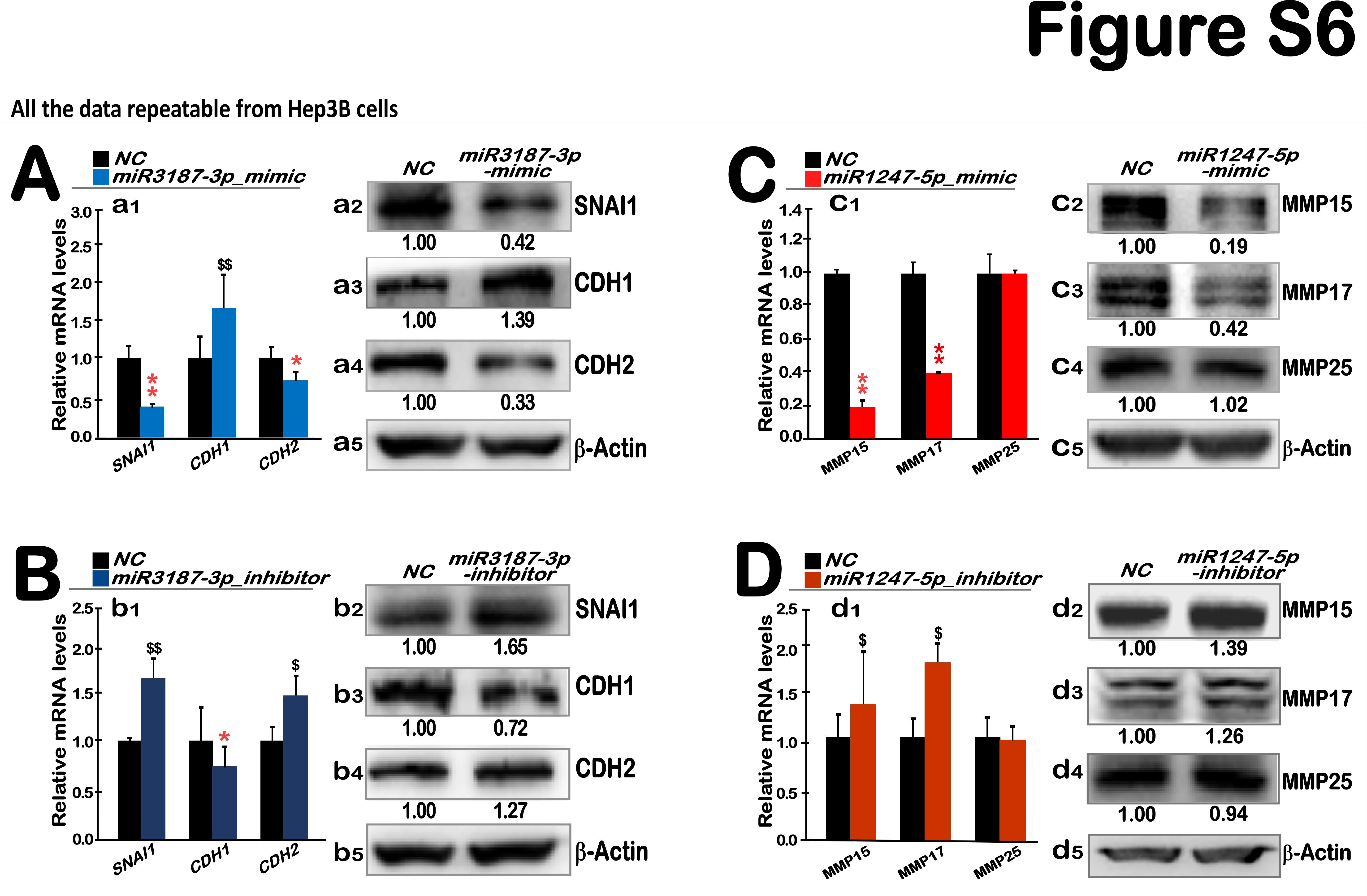

### FigureS7

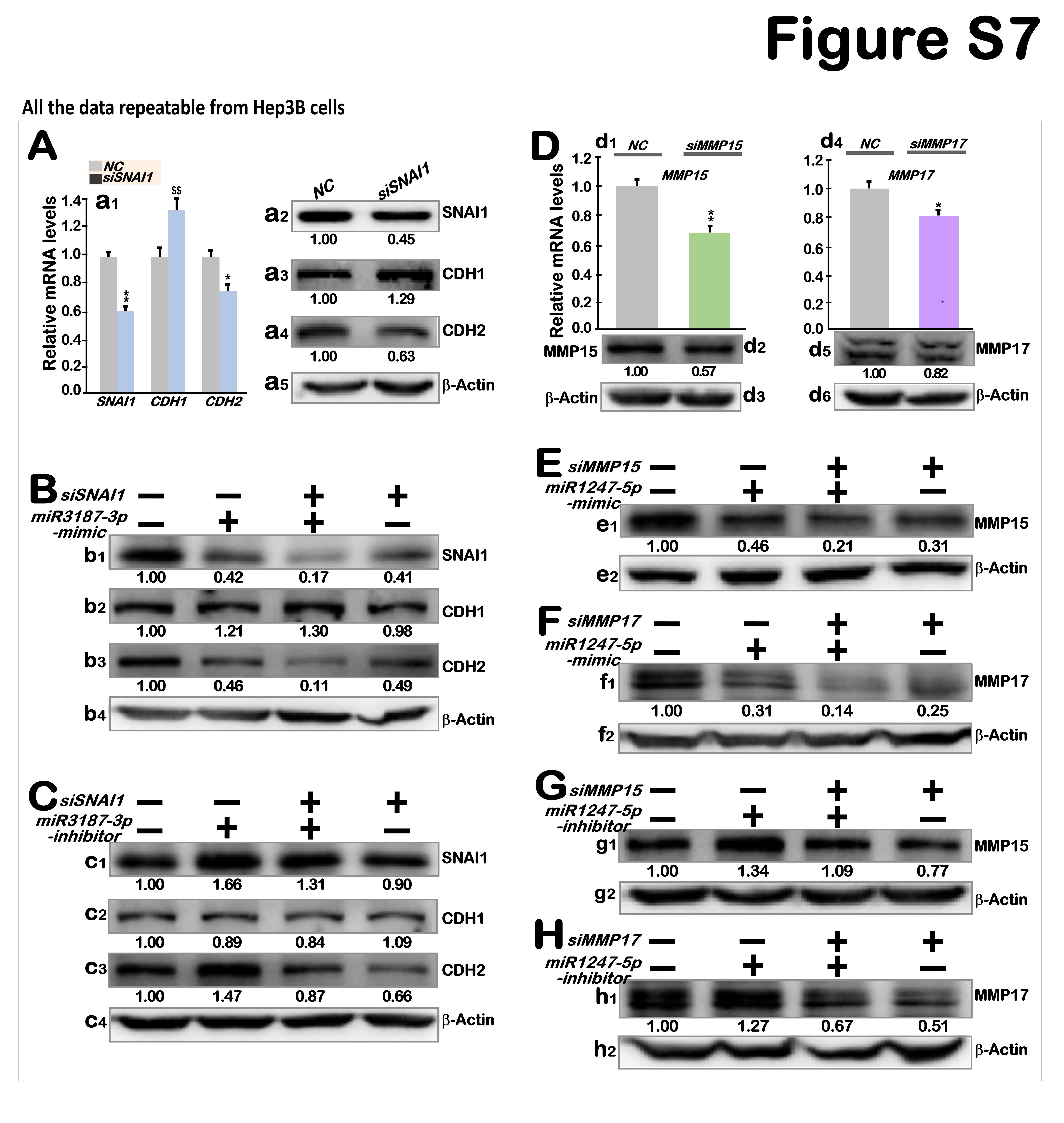

### FigureS8

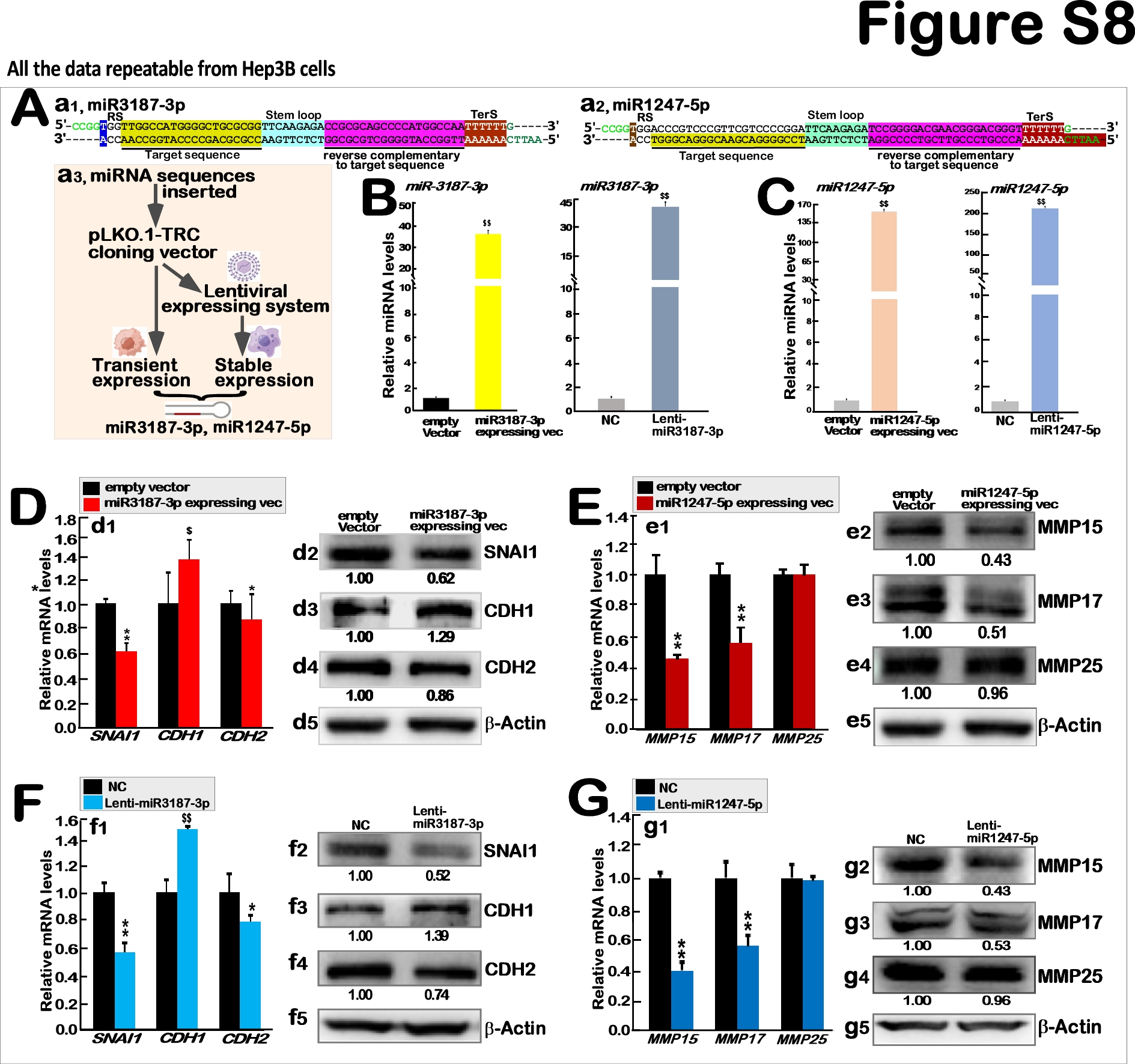

### FigureS9

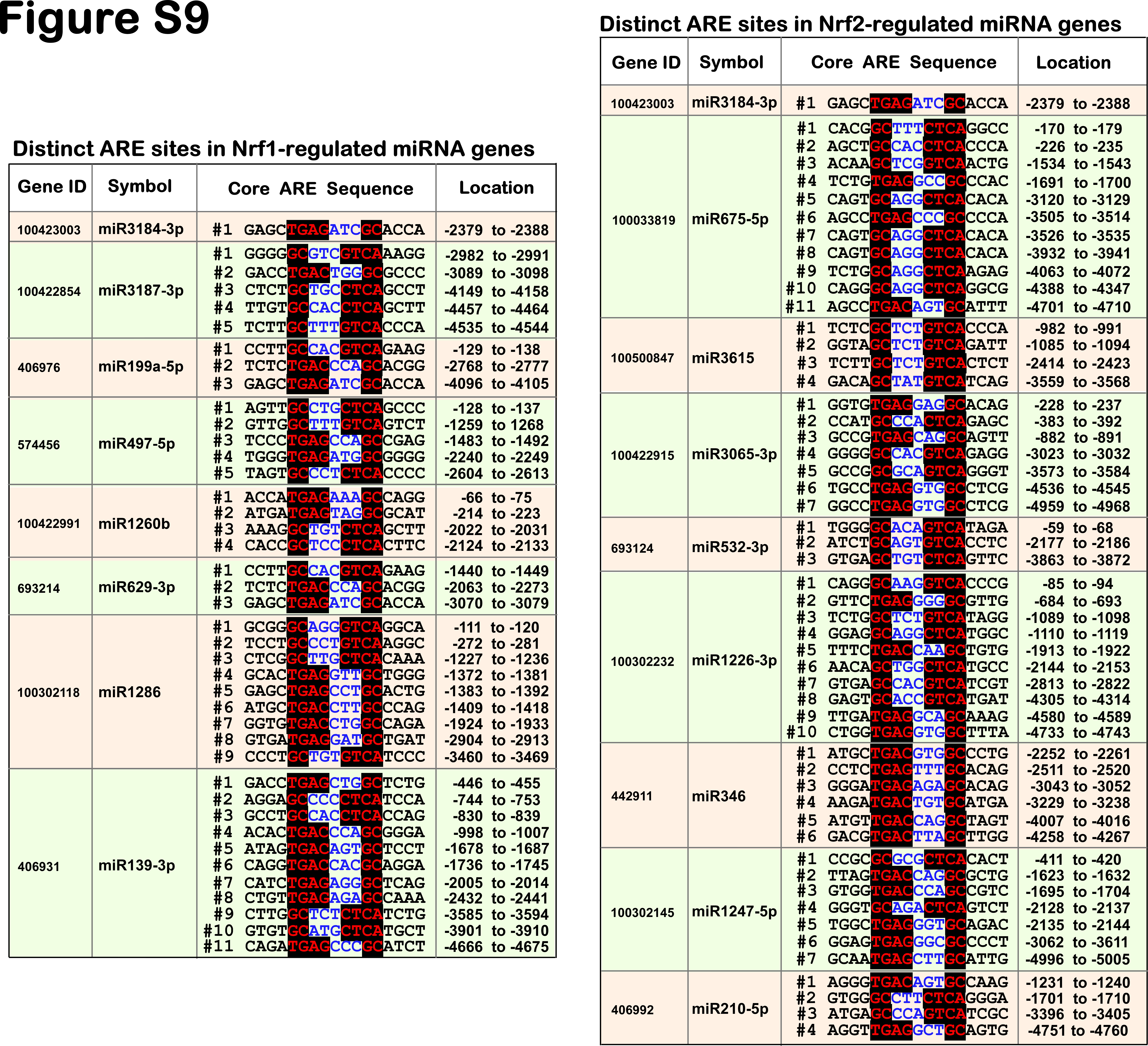

### FigureS10

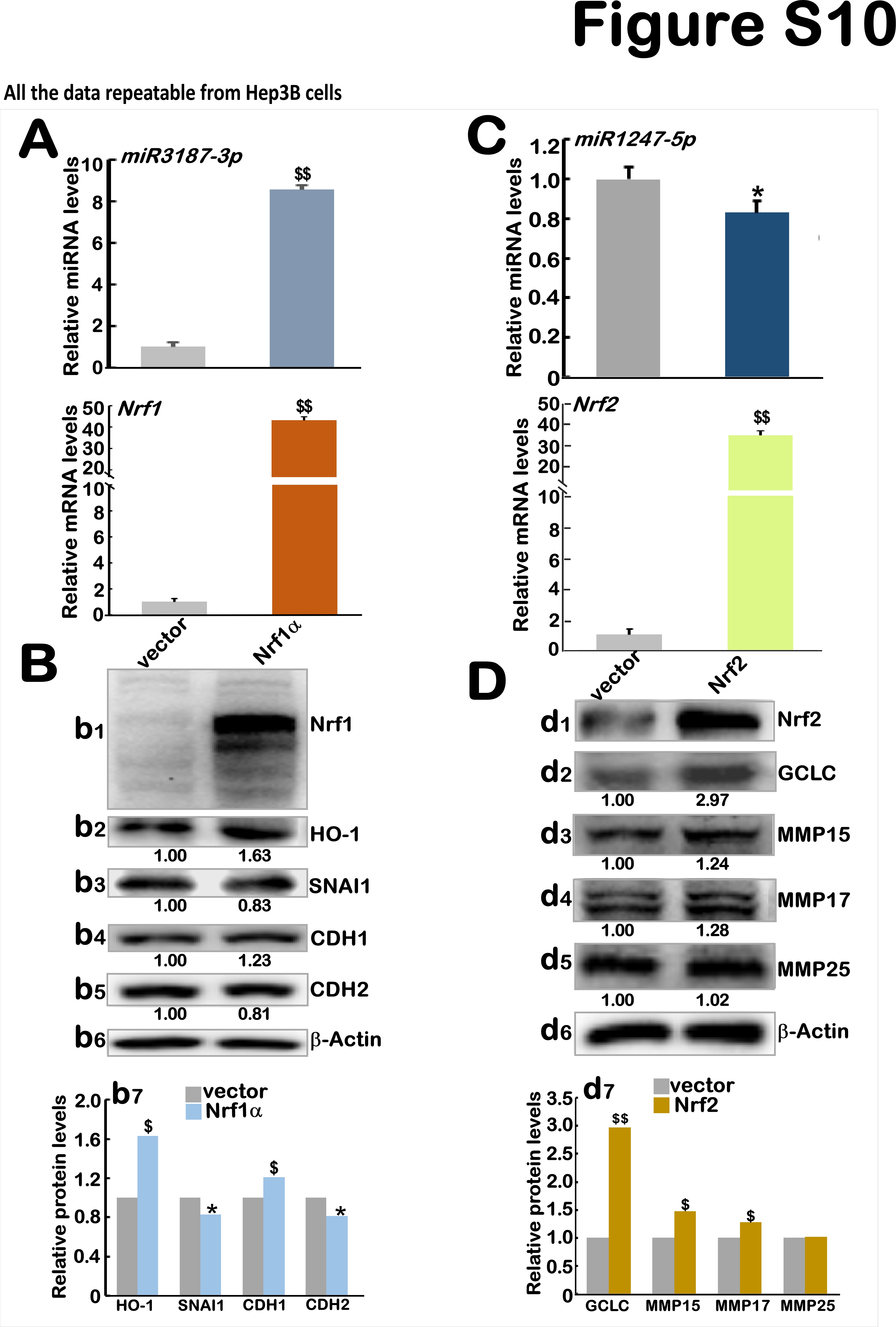

### FigureS11

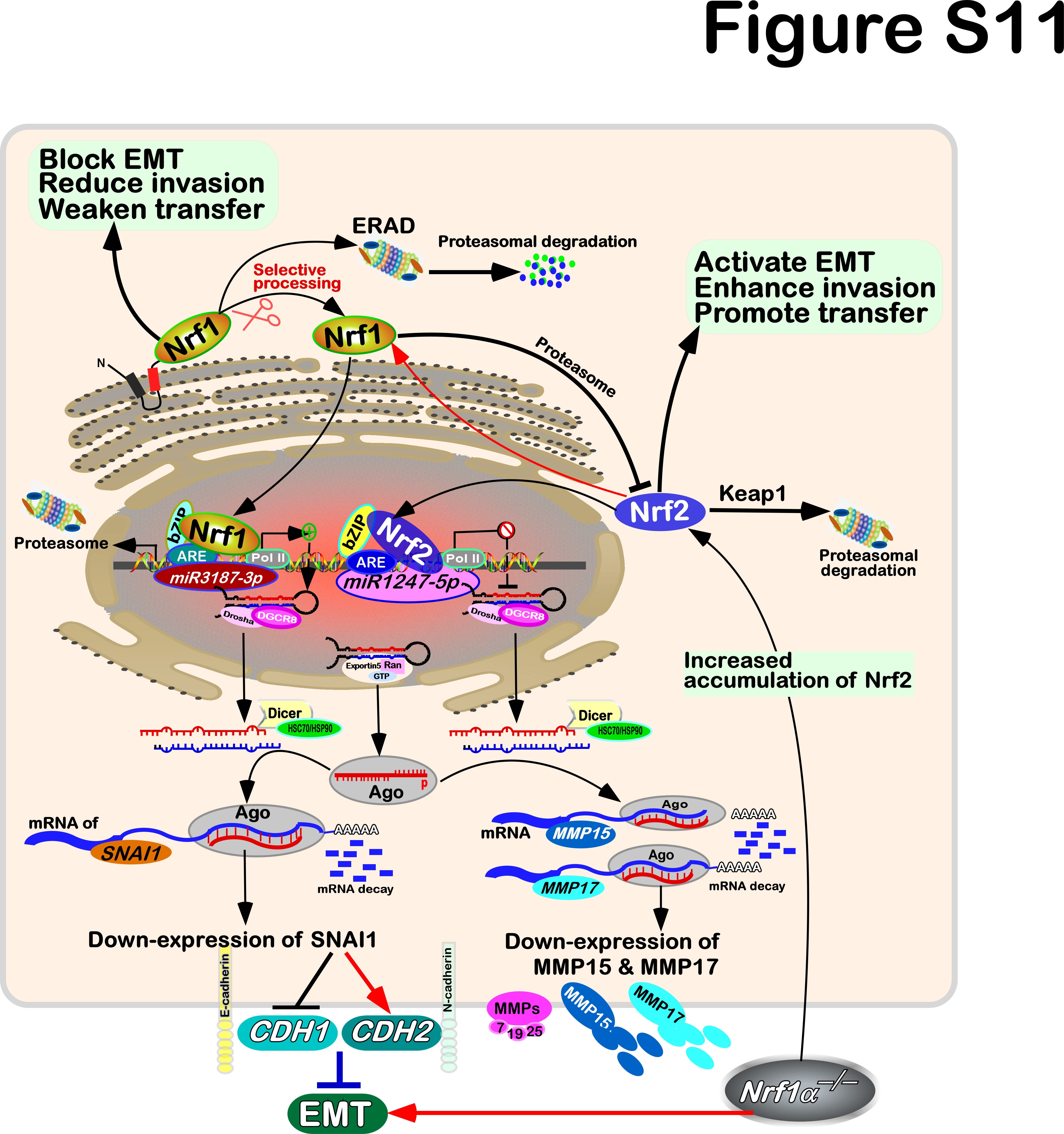

### sFigureS1

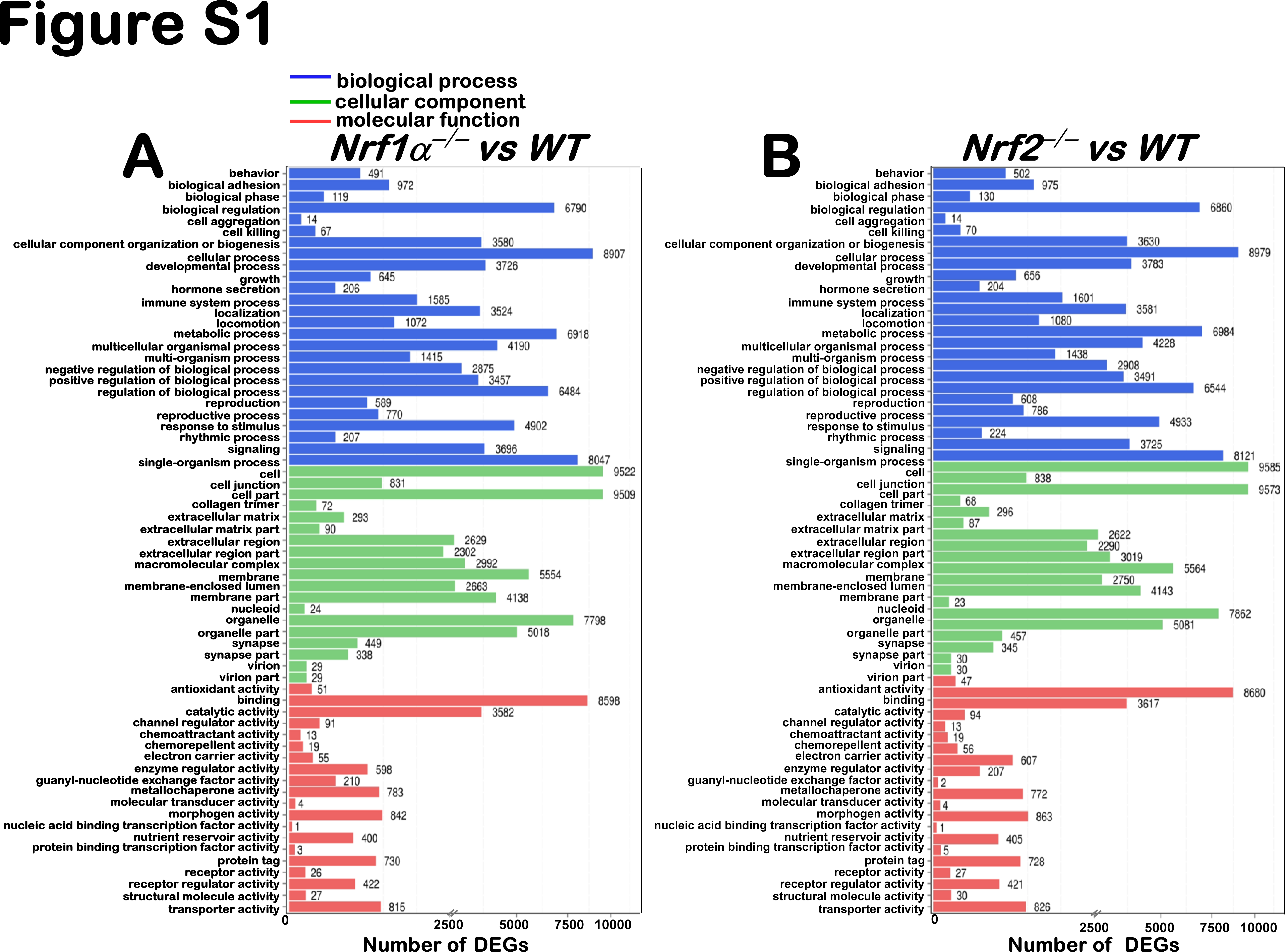
