## Supplementary material for "The opposing mechanisms by which miRNAs critically contribute to differential roles of Nrf1 and Nrf2 in modulating the epithelial-mesenchymal transformation of hepatocellular carcinoma": Table S1 Primers for qPCR

| **Primer name** | **Sequence** | **Corporation** |
| --- | --- | --- |
| 3184-3pRT | GTCGTATCCAGTGCGTGTCGTGGAGTCGGCAATTGCACTGGATACGACTGAGGGG | Tsingke |
| 1286- RT | GTCGTATCCAGTGCGTGTCGTGGAGTCGGCAATTGCACTGGATACGACAGGGCTC | Tsingke |
| 3187-3p-RT | GTCGTATCCAGTGCGTGTCGTGGAGTCGGCAATTGCACTGGATACGACCCGCGCAG | Tsingke |
| 199a-5p-RT | GTCGTATCCAGTGCGTGTCGTGGAGTCGGCAATTGCACTGGATACGACGAACAGG | Tsingke |
| 497-5pRT | GTCGTATCCAGTGCGTGTCGTGGAGTCGGCAATTGCACTGGATACGACACAAACC | Tsingke |
| 629-3pRT | GTCGTATCCAGTGCGTGTCGTGGAGTCGGCAATTGCACTGGATACGACGCTGGGC | Tsingke |
| 1260-bRT | GTCGTATCCAGTGCGTGTCGTGGAGTCGGCAATTGCACTGGATACGACATGGTGG | Tsingke |
| 675-5pRT | GTCGTATCCAGTGCGTGTCGTGGAGTCGGCAATTGCACTGGATACGACCTCTGTG | Tsingke |
| 3615 RT | GTCGTATCCAGTGCGTGTCGTGGAGTCGGCAATTGCACTGGATACGACGAGCCGC | Tsingke |
| 3065-3pRT | GTCGTATCCAGTGCGTGTCGTGGAGTCGGCAATTGCACTGGATACGACTCCAACA | Tsingke |
| 532-3pRT | GTCGTATCCAGTGCGTGTCGTGGAGTCGGCAATTGCACTGGATACGACTGCAAGC | Tsingke |
| 1226-3pRT | GTCGTATCCAGTGCGTGTCGTGGAGTCGGCAATTGCACTGGATACGACCTAGGGA | Tsingke |
| 346RT | GTCGTATCCAGTGCGTGTCGTGGAGTCGGCAATTGCACTGGATACGACAGAGGCA | Tsingke |
| 1247-5pRT | GTCGTATCCAGTGCGTGTCGTGGAGTCGGCAATTGCACTGGATACGACTCCGGGG | Tsingke |
| 210-5pRT | GTCGTATCCAGTGCGTGTCGTGGAGTCGGCAATTGCACTGGATACGACCAGTGTG | Tsingke |
| 139-3pRT | GTCGTATCCAGTGCGTGTCGTGGAGTCGGCAATTGCACTGGATACGACACTCCAA | Tsingke |
| 3184-3p quantitative primer | AAAGTCTCGCTCTCTGCCCCTCA | Tsingke |
| 1286  quantitative primer | TGCAGGACCAAGATGAGCCCT | Tsingke |
| 3187-3p  quantitative primer | ATATTTGGCCATGGGGCTGCGC | Tsingke |
| 199a-5p  quantitative primer | GCGCCCAGTGTTCAGACTACCTG | Tsingke |
| 497-5p  quantitative primer | GCCAGCAGCACACTGTGGTTTG | Tsingke |
| 629-P   quantitative primer | GTTCTCCCAACGTAAGCCCAGC | Tsingke |
| 1260-b   quantitative primer | GCATCCCACCACTGCCACCAT | Tsingke |
| 675-5p  quantitative primer | AAGGAGCGGAGAGGGCCCACAGA | Tsingke |
| 3615  quantitative primer | ATTCTCTCGGCTCCTCGCGGCT | Tsingke |
| 3065-3p  quantitative primer | GCGTCAGCACCAGGATATTGTTGG | Tsingke |
| 532-3p  quantitative primer | ATCCTCCCACACCCAAGGCTTGC | Tsingke |
| 1226-3p  quantitative primer | TCACCAGCCCTGTGTTCCCTAG | Tsingke |
| 346  quantitative primer | TGTCTGCCCGCATGCCTGCCTC | Tsingke |
| 1247-5p  quantitative primer | ATAACCCGTCCCGTTCGTCCCC | Tsingke |
| 210-5p  quantitative primer | ATAGCCCCTGCCCACCGCACACT | Tsingke |
| 139-3p  quantitative primer | TGGAGACGCGGCCCTGTTGGAGT | Tsingke |
| miRNA universal primer R | CGTATCCAGTGCGTGTCGTG | Tsingke |
| miRNA internal reference U6 set F | TCGCTTCGGCAGCACATATACTA | Tsingke |
| miRNA internal reference U6 set R | ATATGGAACGCTTCACGAATTTGC | Tsingke |
| Nrf1 FW | AACATTCTGGTCCTTCAGCAATGCT | Tsingke |
| Nrf1 REV | ACCCGTACCCCAATCAAACTCAGCA | Tsingke |
| CDH1 FW | CAATCCCGATGAAATTGGAA | Tsingke |
| CDH1 REV | TTCATAGTCAAACACGAGCA | Tsingke |
| CDH2 FW | CGCTTTTGTTACATTGCAT | Tsingke |
| CDH2 REV | CAAGATAATAAAATCGCTCCA | Tsingke |
| SNAI1 FW | ATTTCAGCCTCCTGTTTGGT | Tsingke |
| SNAI1 REV | AGCCATTACTCACAGTCCCT | Tsingke |
| MMP15 FW | GTACCCCAAGCCCATCAGT | Tsingke |
| MMP15 REV | TTCCAGTATTTGGTGCCCTT | Tsingke |
| MMP17 FW | CCAGCGACCACAAGATCGTC | Tsingke |
| MMP17 REV | CGGGTATCCTTCCTCTACGTT | Tsingke |
| MMP25 FW | CCCTTCTATCCCCAAGAACCC | Tsingke |
| MMP25 REV | TCTTTCCACCAGCGTTGAGT | Tsingke |
| Twist1 FW | GGCCAGGTACATCGACTTCC | Tsingke |
| Twist1 REV | CTCCATCCTCCAGACCGAGA | Tsingke |
| TP53 FW | ACCTACCTCACAGAGTGCAT | Tsingke |
| TP53 REV | ACTACCAACCCACCGACCAA | Tsingke |
| FN1 FW | TCCGAGTCAGCCCAACTCC | Tsingke |
| FN1 REV | AGCTTCCTTCCAACGGCCTA | Tsingke |
| MMP2 FW | CCGATAACCTGGATGCCGTC | Tsingke |
| MMP2 REV | CTCAGCAGCCTAGCCAGTCG | Tsingke |
| MMP11 FW | GGGGTACAACCACCATGACA | Tsingke |
| MMP11 REV | GCCCTTCCTCTTAAGTCATGC | Tsingke |
| MMP16 FW | ATGCCTAAAACCTTCATCCTT | Tsingke |
| MMP16 REV | AAGCTGCCTCTGTAACACC | Tsingke |
| MMP19 FW | CAACTGGATGCACTGTCGTC | Tsingke |
| MMP19 REV | CAAGGTTATGCCCGTACCTG | Tsingke |
| MMP24 FW | ACCGTCACCTCCTTAAACACC | Tsingke |
| MMP24 REV | CCCTTCTATCCCCAAGAACCC | Tsingke |
| MMP25 FW | CCCTTCTATCCCCAAGAACCC | Tsingke |
| MMP2 FW | TCTTTCCACCAGCGTTGAGT | Tsingke |
| MMP27 FW | CAAGCTGCATACGAGAACCC | Tsingke |
| MMP27 FW | CTGGCAAGACAGCATATCCTC | Tsingke |
