## Supplementary material for "The opposing mechanisms by which miRNAs critically contribute to differential roles of Nrf1 and Nrf2 in modulating the epithelial-mesenchymal transformation of hepatocellular carcinoma": Table S2 Primers for plasmid construction

| **Primer name** | **Sequence** | | **Corporation** | |
| --- | --- | --- | --- | --- |
| Whole length 3187F： | GGGTACCCTCCAGGCTCTTCTGCTTCCACTCGC | | Tsingke | |
| Whole length 3187R: | CGGAATTCCGCGGCAAGTCCAACGATCACAG | | Tsingke | |
| Whole length 1247F: | GGGGTACC GCCCCGTCCCATCTCGTTCCAGC | | Tsingke | |
| Whole length 1247R: | CGGAATTC CCCTGAGCCCAAGTCGCCAACCC | | Tsingke | |
| Mutation SNAIlF: | GCTCTGGAAGAGGCCTTCCCCGAACTGTTTCTGTGGAGGGAGGGCAG | | Tsingke | |
| Mutation SNAI1R: | CTGCCCTCCCTCCACAGAAACAGTTCGGGGAAGGCCTCTTCCAGAGC | | Tsingke | |
| Mutation MMP17F: | GCTCTGCTGGCGCGGATGATTGAGAAC CACCCTGGCACCGGAAGG | | Tsingke | |
| Mutation MMP17R: | CCTTCCGGTGCCAGGGTGGCCCGTCCTCATCCGCGCCAGCAGAGC | | Tsingke | |
| Mutation MMP15F: | GCTCTTGGGTTCTGAGCACTTACTTGAGGAAGTGGAACAACCA | | Tsingke | |
| Mutation MMP15R: | CTGCCACCCAAGACTCGTGAATGAACTCCTTCACCTTDTTGGT | | Tsingke | |
| Mutation MMP25F: | GCTCTCTTGTCTTCCCACTGCATGA CACCCTGGCACCGGAAGG | | Tsingke | |
| Mutation MMP25R: | CCTTCCGGTGCCAGGGTGTCATGCAGTGGGAAGACAAGAGAGC | | Tsingke | |
| Luci-3187-1F: | GGTACCGGGGCGTCGTCAAAGGGGGCGTCGTCAAAG GGGGCGTCGTCAAAGC | | Tsingke | |
| Luci-3187-1R: | TCGAGGCTTTGACGACGCCCCCTTTGACGACGCCCCCTTTGACGACGCCCCG | | Tsingke | |
| Luci-3187-2F: | GGTACCACCTGACTGGGCGCCACCTGACTGGGCGCC ACCTGACTGGGCGCCC | | Tsingke | |
| Luci-3187-2R: | TCGAGGGCGCCCAGTCAGGTGGCGCCCAGTCAGGTGGCGCCCAGTCAGGTG | | Tsingke | |
| Luci-3187-3F: | GGTACCTCTGCTGCCTCAGCCTCTGCTGCCTCAGCC TCTGCTGCCTCAGCCC | | Tsingke | |
| Luci-3187-3R: | | TCGAGGGGCTGAGGCAGCAGAGGCTGAGGCAGCAGAGGCTGAGGCAGCAGAG | | Tsingke |
| Luci-3187-4F: | | GGTACCTGTGCCACCTCAGCTTGTGCCACCTCAGCT TGTGCCACCTCAGCT | | Tsingke |
| Luci-3187-4R: | | TCGAGGAGCTGAGGTGGCACAAGCTGAGGTGGCACAAGCTGAGGTGGCACAG | | Tsingke |
| Luci-3187-5F: | | GGTACCCTTGCTTTGTCACCC CTTGCTTTGTCACCC CTTGCTTTGTCACCCC | | Tsingke |
| Luci-3187-5R: | | TCGAGGGGGTGACAAAGCAAGGGGTGACAAAGCAAGGGGTGACAAAGCAAG G | | Tsingke |
| Luci-1247-1F: | | GGTACCCGCGCGCGCTCACACCGCGCGCGCTCACAC CGCGCGCGCTCACACC | | Tsingke |
| Luci-1247-1R: | | TCGAGGGTGTGAGCGCGCGCGGTGTGAGCGCGCGCGGTGTGAGCGCGCGCGG | | Tsingke |
| Luci-1247-2F: | | GGTACCTAGTGACCAGGCGCTTAGTGACCAGGCGCT TAGTGACCAGGCGCTC | | Tsingke |
| Luci-1247-2R: | | TCGAGGTAGTGACCAGGCGCT TAGTGACCAGGCGCT TAGTGACCAGGCGCTG | | Tsingke |
| Luci-1247-3F: | | GGTACCTGGTGACCCAGCCGT TGGTGACCCAGCCGT TGGTGACCCAGCCGTC | | Tsingke |
| Luci-1247-3R: | | TCGAGGACGGCTGGGTCACCAACGGCTGGGTCACCAACGGCTGGGTCACCA | | Tsingke |
| Luci-1247-4F: | | GGTACCGGGTGCAGACTCAGTCGGGTGCAGACTCAGTC GGGTGCAGACTCAGTCC | | Tsingke |
| Luci-1247-4R: | | TCGAGGGACTGAGTCTGCACCCGACTGAGTCTGCACCCGACTGAGTCTGCACCCG | | Tsingke |
| Luci-1247-5F: | | GGTACCGGCTGAGGGTGCAGAGGCTGAGGGTGCAGA GGCTGAGGGTGCAGAC | | Tsingke |
| Luci-1247-5R: | | TCGAGGTCTGCACCCTCAGCCTCTGCACCCTCAGCCTCTGCACCCTCAGCC G | | Tsingke |
| Luci-1247-6F: | | GGTACCGAGTGAGGGCGCCCCGAGTGAGGGCGCCCC GAGTGAGGGCGCCCCC | | Tsingke |
| Luci-1247-6R: | | TCGAGGGGGGCGCCCTCACTCGGGGCGCCCTCACTCGGGGCGCCCTCACTCG | | Tsingke |
| Luci-1247-7F: | | GGTACCCAATGAGCTTGCATTCAATGAGCTTGCATT CAATGAGCTTGCATTC | | Tsingke |
| Luci-1247-7R: | | TCGAGGAATGCAAGCTCATTGAATGCAAGCTCATTGAATGCAAGCTCATTGG | | Tsingke |
