## Supplementary material for "The opposing mechanisms by which miRNAs critically contribute to differential roles of Nrf1 and Nrf2 in modulating the epithelial-mesenchymal transformation of hepatocellular carcinoma": Table S3 Information on the antibodies

| Antibodies | Reagent number | Manufacturer |
| --- | --- | --- |
| Nrf1 | Made in our laboratory | BIMAKE，USA |
| Nrf2 | A5448 | BIMAKE，USA |
| SNAI1 | ab108427 | Abcam，UK |
| SNAI2 | AF3418 | R&D Systems |
| MMP15 | bs1855R | BIOSS，China |
| MMP17 | A3030 | Abcam，UK |
| MMP25 | A3032 | Abcam，UK |
| GCLC | D123963 | Sangon Biotechnology，China |
| HO1 | ab52947 | Abcam，UK |
| CDH1 | bs-10009R | BIOSS，China |
| CDH2 | AO1377A | ABGENT，China |
| FN1 | BA1771 | BOSTER，China |
| ITGB4 | BA4486 | BOSTER，China |
| MMP9 | bs-4593R | BIOSS，China |
