## Supplementary material for "The opposing mechanisms by which miRNAs critically contribute to differential roles of Nrf1 and Nrf2 in modulating the epithelial-mesenchymal transformation of hepatocellular carcinoma": Table S4 The cell lines utilized in this study

| Cell line name | Cell lineage species | Cell line source |
| --- | --- | --- |
| HepG2 | Human Hepatocellular Carcinoma | Maintained in our laboratory |
| Hep3B | Human Hepatocellular Carcinoma | Maintained in our laboratory |
| Nrf1^–∕–^ cells | Knockdown of Nrf1 in HepG2 cells | Maintained in our laboratory |
| Nrf2^–∕–^ cells | Knockdown of Nrf2 in HepG2 cells | Maintained in our laboratory |
| 3187-vector | miR-3187-3p was constructed into pLKO.1-TRC cloning vector plasmid | Constructed in our laboratory |
| 1247-vector | miR-1247-5p was constructed into pLKO.1-TRC cloning vector plasmid | Constructed in our laboratory |
| G2-Lentiv-NC | HepG2 cells packed with miRNA-NC | Constructed in our laboratory |
| 3B-Lentiv-NC | Hep3B cells packed with miRNA-NC | Constructed in our laboratory |
| G2-Lentiv-1247 | Stable overexpression of miR-1247-5p in HepG2 cells | Constructed in our laboratory |
| G2-Lentiv-3187 | Stable overexpression of miR-3187-3p in HepG2 cells | Constructed in our laboratory |
| 3B-Lentiv-1247 | Stable overexpression of miR-1247-5p in Hep3B cells | Constructed in our laboratory |
| 3B-Lentiv-3187 | Stable overexpression of miR-3187-3p in Hep3B cells | Constructed in our laboratory |
