## Supplementary material for "The opposing mechanisms by which miRNAs critically contribute to differential roles of Nrf1 and Nrf2 in modulating the epithelial-mesenchymal transformation of hepatocellular carcinoma": Table S5 Glossary of abbreviations

| Abbreviation | English full title |
| --- | --- |
| AKT | Protein kinase B |
| CDH1 | cadherin 1 |
| CDH2 | cadherin 2 |
| c-Myc | MYC Proto-Oncogene |
| ECM | Extracellular matrix |
| EGF | Epidermal growth factor |
| EMT | Epithelial-Mesenchymal Transition |
| EMT-TFs | EMT-related transcription factors |
| FN1 | Fibronectin 1 |
| GCLC | Glutamate-Cysteine Ligase Catalytic Subunit |
| HCC | Hepatocellular carcinoma |
| miRNAs | MicroRNAs |
| MMPs | Matrix metallopeptidase families |
| MMP11/15/17/25 | Matrix Metallopeptidase 11/15/17/25 |
| MAPK | Mitogen-activated protein kinase |
| Nrf1/2 | Nuclear Respiratory Factor1/2 |
| Prrx1 | Transcription factor pairing-related homeobox 1 |
| PI3K | phosphatidylinositol-4,5-bisphosphate 3-kinase |
| PTEN | phosphatase and tensin homolog |
| SNAI 1/2 | Snail Family Transcriptional Repressor 1/2 |
| TTF1 | thyroid transcription factor 1 |
| TGF-β | Transforming growth factor-β |
| UTR | Untranslated region |
| VIM | Vimentin |
| WB | Western blotting |
| WNT1 | Wnt family member 1 |
